# Identification of reactive CpGs and RNA expression in early COVID-19 through *cis*-eQTM analysis reflecting disease severity and recovery

**DOI:** 10.1101/2025.06.09.658549

**Authors:** Hyojung Ryu, Kyungwhan An, Yoonsung Kwon, Yeonsu Jeon, Sungwon Jeon, Hansol Choi, Yeo Jin Kim, Sunhwa Kim, Ok Joo Sul, SangJoon Lee, Asaph Young Chun, Eun-Seok Shin, Seung Won Ra, Jong Bhak

**Affiliations:** Clinomics, Inc., Ulsan, 44919, Republic of Korea; GenomeLab, Korean Genomics Center (KOGIC), Ulsan National Institute of Science and Technology (UNIST), Ulsan, 44919, Republic of Korea; Department of Biomedical Engineering, College of Information and Biotechnology, Ulsan National Institute of Science and Technology (UNIST), Ulsan, 44919, Republic of Korea; Geromics Inc., Suwon, 16229, Republic of Korea; AgingLab, Ulsan, 44919, Republic of Korea; Division of Pulmonology, Department of Internal Medicine, Ulsan University Hospital, University of Ulsan College of Medicine, Ulsan, Republic of Korea; Biomedical Research Center, Ulsan University Hospital, School of Medicine, University of Ulsan, Ulsan, 44033, Republic of Korea; Basic-Clinical Convergence Research Institute, University of Ulsan, Ulsan, 44610, Republic of Korea; Department of Biological Science, Ulsan National Institute of Science and Technology (UNIST), Ulsan, Republic of Korea; Institute for Pandemic Sciences AI.celerator, Seoul National University, Seoul, 08826, Republic of Korea; Department of Cardiology, Ulsan University Hospital, University of Ulsan College of Medicine, Ulsan, Republic of Korea

## Abstract

Multi-omics analyses of severe COVID-19 cases are crucial in deciphering the complex interplay between genetic and epigenetic factors. Here, we present an analysis of Expression Quantitative Trait Methylation (eQTM) to investigate the complex interplay of methylation and gene expression pattern during the acute phase of severe COVID-19. We identified 16 differentially expressed genes and 30 nearby differentially methylated CpG sites. Six key genes—*SRXN1*, *FURIN*, *IL18RAP*, *FOXO3*, *GCNT4*, and *FKBP5*—were either up-regulated or down-regulated near hypomethylated CpG sites. These genes are associated with viral infiltration, immune activation, lung damage, and oxidative stress-related multi-organ failure, which are the hallmarks of severe COVID-19. Interestingly, during the recovery phase, methylation and gene expression levels returned to baseline, underscoring the rapid and reversible nature of these molecular changes. These findings provide insight into the dynamics of epigenetic and transcriptomic shifts according to the infectious stage, supporting potential prognostic and therapeutic approaches for severe COVID-19.

## Introduction

The global impact of the coronavirus disease 2019 (COVID-19) has underscored the urgent need to understand the more precise molecular mechanisms underlying its clinical manifestations, particularly the factors contributing to severe disease outcomes^1^. The clinical spectrum of COVID-19 ranges from asymptomatic cases to severe respiratory failure, yet the underlying molecular and omics drivers of these varied outcomes remain elusive. COVID-19 can trigger a cytokine storm—an excessive immune response characterized by the overproduction of cytokines—that results in tissue damage and widespread inflammation, which are strongly linked to severe disease^2^. In the most critical cases, the cytokine storm spreads to multiple organs, ultimately causing multi-organ failure and death^3^.

In recent years, big-data-driven precision multi-omics approaches have emerged as a powerful tool for dissecting the complex biological interactions that underpin pathologies. By integrating multiple layers of omics data, such as genomics, transcriptomics, and epigenomics, these precision medicine approaches can be utilized to gain a comprehensive understanding of disease mechanisms in instances such as COVID-19. DNA methylation, as the most common epigenetic modification, plays a crucial role in regulating gene expression and can be influenced by infections, as well as other stressors^4,5^. Several studies have demonstrated that epigenetic regulation plays a role in the severity of COVID-19^4,6–9^. Previously, DNA methylation was considered relatively stable than other epigenetic modifications. However, recent studies have revealed that DNA methylation can occur more rapidly than previously thought^10^. This is particularly evident when cells are exposed to challenging environments such as infection caused by direct invasion of pathogens. RNA expression analysis is the most informative molecular method measuring the effect and severity of infectious diseases such as COVID-19, because it directly and rapidly reflects the epigenetic changes caused by viral infection^11^. Therefore, Expression Quantitative Trait Methylation (eQTM) analysis has been successfully employed in several studies to investigate the relationship between DNA methylation and gene expression^12–14^. However, despite its potential, eQTM approaches have rarely been applied to investigate the connection between DNA methylation and RNA expression in relation to COVID-19 severity.

In this study, we employed a *cis-*eQTM analysis to investigate the integrated patterns of DNA methylation and RNA expression in 46 patients with varying degrees of COVID-19 severity. Our objective was to determine the molecular mechanisms associated with severe outcomes during the acute phase of the disease and to assess the dynamic changes in multi-omics of these molecular signatures during the recovery phase. These observations underscore the dynamic nature of epigenetic and transcriptomic changes in COVID-19 progression and recovery, providing valuable insights into potential healthcare and therapeutic targets for mitigating severe outcomes.

## Results

### Dynamics of gene-regulating CpGs and gene expression levels in COVID-19 severity

We analyzed blood-based multi-omic differences between severe-critical (SC) and mild-moderate (MM) cases during the acute phase of infection in 46 patients hospitalized with COVID-19 (Figure 1A). The clinical follow-up period for all patients is presented in Figure 1B. We confirmed their severity status based on clinical lab values, including blood cell counts and inflammatory biomarker levels. Notably, SARS-CoV-2 viral load, as measured by PCR cycle threshold (Ct) values for multiple gene targets (*N*, *E*, and *R* genes), did not differ significantly between severity groups (Figure S1), suggesting that differences in clinical outcomes were not attributable to viral burden alone but to host-intrinsic factors, including immune and epigenetic responses.

**Figure 1.**
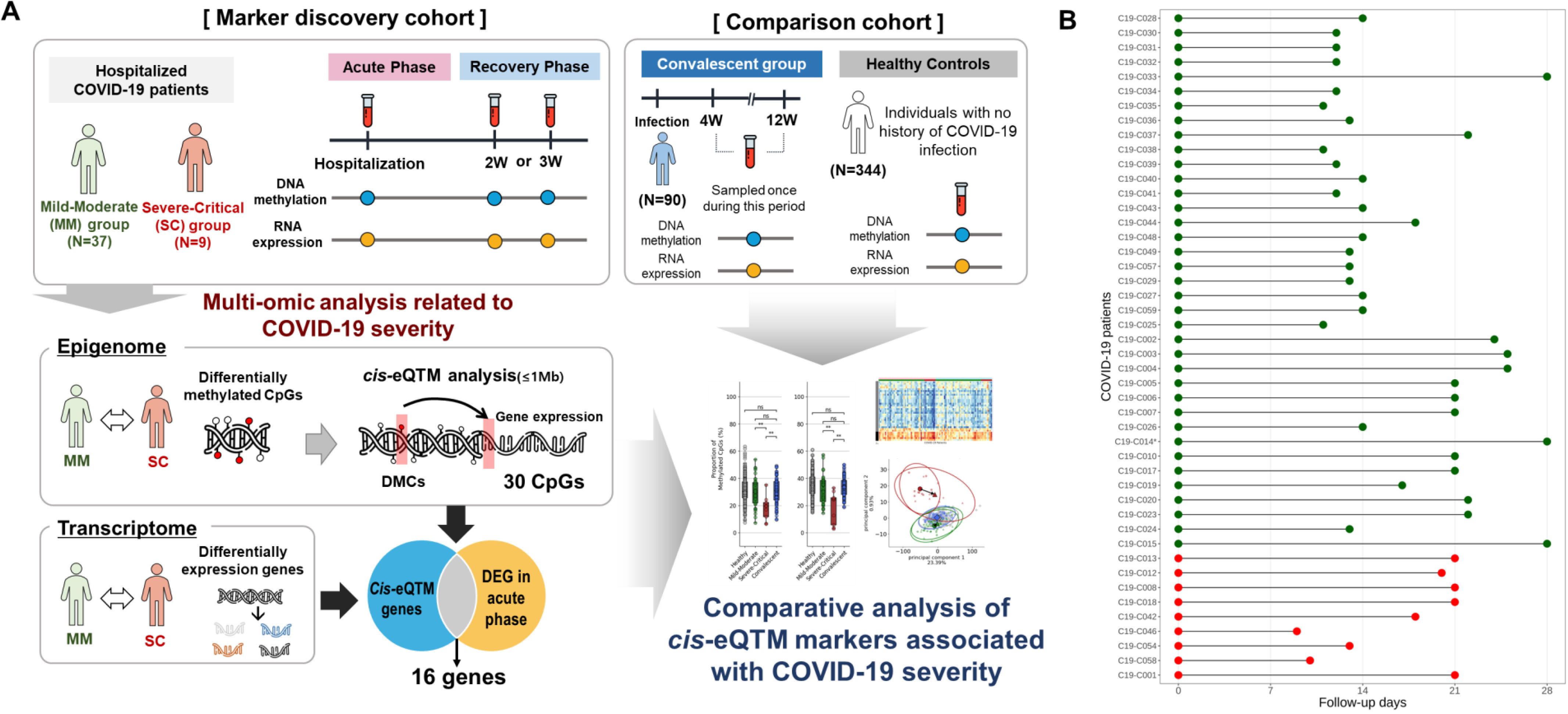
Overall study design and follow-up blood sample collection for *cis-*eQTM analysis in COVID-19 patients. **A)** Overview of blood sampling and analysis design; The marker discovery cohort consisted of hospitalized COVID-19 patients, stratified into Mild-Moderate (MM) (N=37) and Severe-Critical (SC) (N=9) groups, and was used to identify severity-associated *cis*-eQTM markers. The comparison cohort was composed of two independent reference groups: the ‘Convalescent group’ (N=90) comprising individuals who recovered from COVID-19 and provided a single blood sample 4 to 12 weeks post-infection, and the ‘Healthy Controls’ group (N=344), consisting of pre-pandemic individuals with no history of SARS-CoV-2 infection. **B)** Clinical follow-up duration of 46 Hospitalized COVID-19 patients. The green and red points indicate MM and SC groups, respectively. Sample C19-C014(*), excluded DEG analysis due to QC failure, was retained in methylation analysis.

Leave-one-out (LOO) analysis between SC and MM COVID-19 groups identified 648 hypermethylated and 1,296 hypomethylated CpG sites in the severe disease. *Cis*-eQTM analysis further revealed that 732 of these differentially methylated CpGs (113 hypermethylated CpGs and 619 hypomethylated CpGs) were correlated with 928 genes located within ±1Mbp of the differentially methylated CpGs (DMCs). Transcriptome-wide differential expression analysis identified 297 differentially expressed genes (DEGs) between SC and MM COVID-19 groups that were consistently observed in at least seven LOO iterations. Of these, 283 genes were up-regulated and 14 down-regulated in the severe disease. The up-regulated genes were enriched in pathways related to erythrocyte dynamics, innate immune activation, platelet aggregation, and acetylcholine receptor signaling (Figure S2A), whereas down-regulated genes were predominantly involved in DNA mismatch repair (MMR) pathway (Figure S2B).

Integration of methylation and transcriptomic data through *cis*-eQTM analysis refined these genes to 16 DEGs that were proximal to 30 DMCs (Table S1). Among these, 15 genes were up-regulated and a single gene, *GCNT4*, was down-regulated. The overlap of these DEGs with *cis-*eQTM loci indicates their expression is likely regulated by local methylation status (Table S1).

From the final set of *cis-*eQTM genes, six key genes and their ten regulatory CpGs associated with COVID-19 infection and severity were selected for further analysis based on their previous association with COVID-19 severity (Table 1). These genes are mechanistically linked to viral entry, immune regulation, and oxidative stress–major hallmarks of severe disease. For instance, *FURIN*, *SRXN1*, and *FKBP5*, which were up-regulated in the SC group, had strong negative correlations with their associated CpG methylation levels (ρ = –0.602, – 0.582 to –0.510, and –0.551, respectively; *P* = 0.003, –0.004 to 0.021, and 0.009, respectively), indicating that hypomethylation at these loci is linked to increased gene expression. *FOXO3* and *IL18RAP* also showed moderate inverse correlations (ρ = –0.501 and –0.503; *P* = 0.025 and 0.024), suggesting methylation-dependent regulation. In contrast, *GCNT4*, the only down-regulated gene among the six, was positively correlated with methylation levels (ρ = 0.528; *P* = 0.015).

**Table 1.**
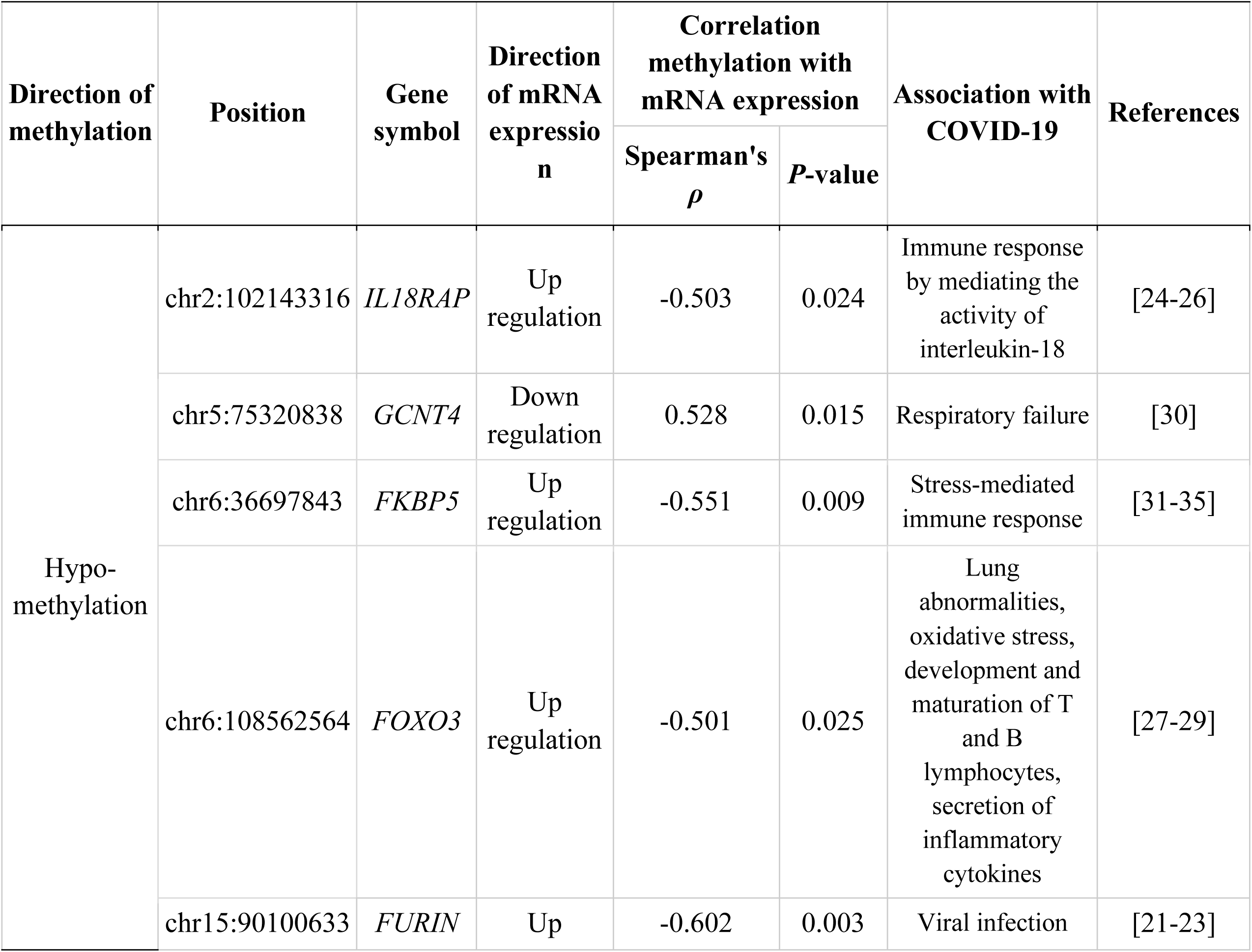
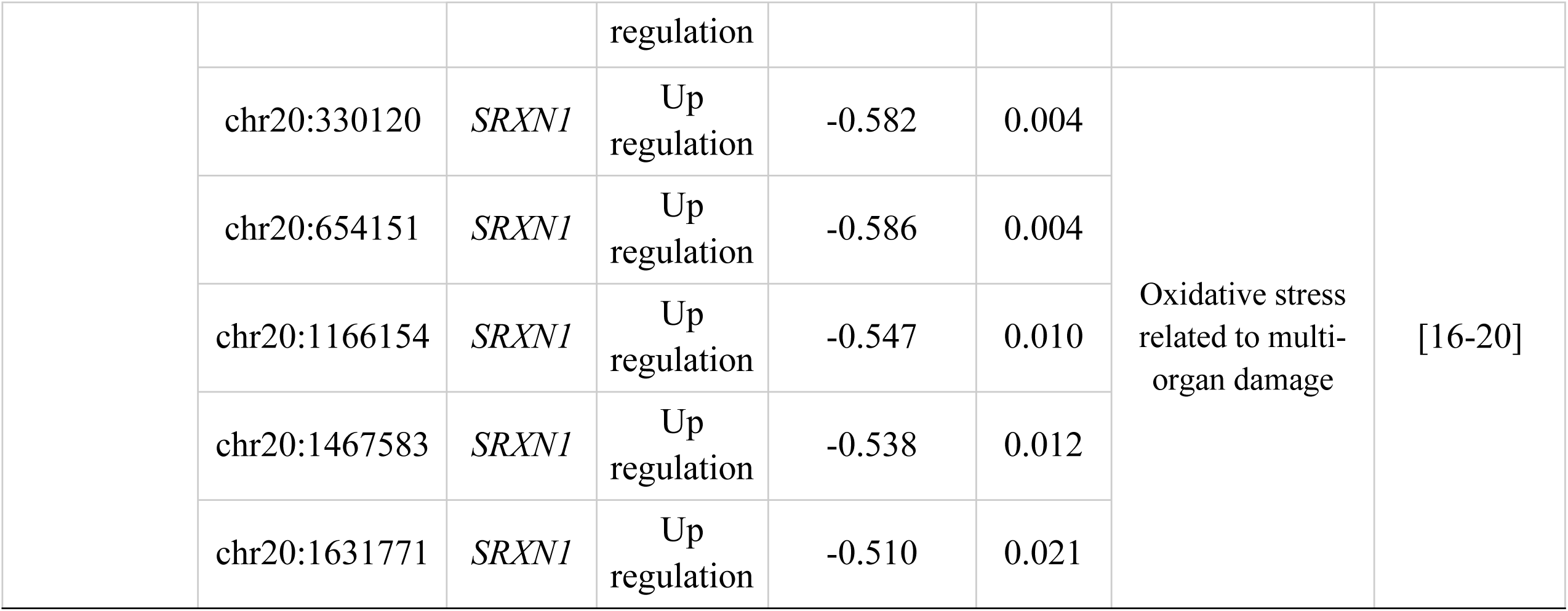
Six key genes associated with the severity of COVID-19 in the acute phase. Columns indicate the direction of DNA methylation change (hypo- or hypermethylation); CpG site genomic position (hg38); associated gene symbol; direction of mRNA expression change (up- or downregulation); direction of correlation between DNA methylation and gene expression, Spearman’s correlation coefficient (ρ), and associated *P*-value from *cis*-eQTM analysis; previously reported associations of each gene with COVID-19 and relevant references.

We examined the distributions of DNA methylation and expression of the six prioritized genes according to the infection phase. Mean methylation differences between MM and SC COVID-19 patients were statistically significant for all ten CpGs (Table S2). The convalescent group, with samples collected from 4 to 12 weeks post-COVID-19 infection, showed no significant differences in methylation levels compared to the healthy control group. Notably, the convalescent individuals from the MM group during the acute phase presented no significant differences in their methylation profiles across nine of the ten CpGs, with the exception of chr20:1166154 which regulates *SRNX1* expression (Figure 2A, Figure S3). The pattern was mirrored at the gene expression level (Figure 2B). All six key genes were either up- or down-regulated in the SC group compared to the MM group. Three of the six genes (*FURIN*, *SRXN1*, and *FKBP5*) displayed significant differences in mean expression only between severe patients compared to convalescent individuals, but not between the MM and convalescent group (Table S3). As for markers with no previous report on COVID-19 severity, all 24 DMCs showed significantly altered methylated proportions (Figure S3). Among ten DEGs with no past association with COVID-19 severity, nine genes (except *GABARAPL2*) displayed significant expression changes between MM and SC (Figure S4).

**Figure 2.**
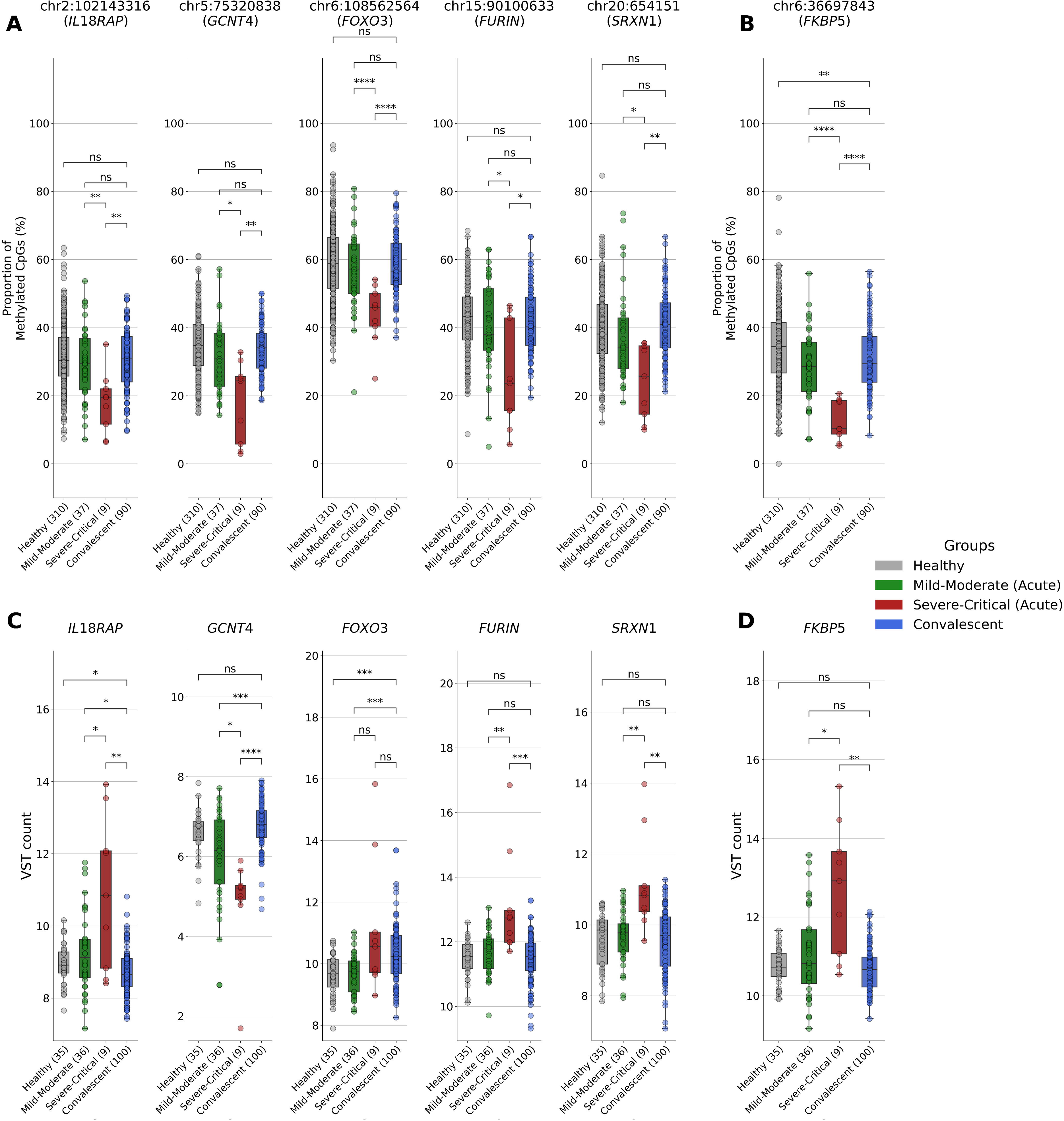
Multiomic Profiling of Six Key *cis*-eQTM Loci Reveals COVID-19 Severity-Associated Methylation and Gene Expression Changes Across the Infection Timeline. **A,C)** DNA methylation levels (% methylated CpGs) at representative cis-eQTM sites, stratified by clinical group: Healthy controls (n=310), Mild-Moderate (acute phase; n=37), Severe-Critical (acute phase; n=9), and Convalescent individuals (n=90). Panel **A** presents five cell-type dependent cis-eQTM CpGs, whereas panel **C** illustrates a cell-type independent CpG. **B, D)** Gene expression levels (VST count) for the matched loci are displayed in panels **B** (cell-type proportion dependent) and **D** (cell-type proportion independent) across clinical group: Healthy controls (n = 35), Mild-Moderate (acute phase; n = 36), Severe-Critical (acute phase; n = 9), and Convalescent individuals (n = 90). Boxplots display the median, interquartile range (IQR), and individual data points. Statistical significance was evaluated using Welch’s *t*-test for methylation and the Wilcoxon rank-sum test for gene expression, with nominal *P*-values. Significance thresholds are indicated as follows: ns (*P* ≥ 0.05), *(*P* < 0.05), ** (*P* < 0.01), *** (*P* < 0.001), **** (*P* < 0.0001). Abbreviation: VST, variance-stabilizing transformation.

### Leukocyte-adjusted *cis*-eQTM analysis reveals *FKBP5* methylation as a cell-type proportion-independent marker of severe COVID-19

Because acute SARS-CoV-2 infection disrupts circulating leukocyte distributions, we repeated the *cis*-eQTM analysis while additionally adjusting for neutrophil, lymphocyte, monocyte, eosinophil, and basophil proportions obtained from clinical blood counts. Consistent with this premise, a direct comparison of the measured cell proportion revealed marked baseline differences between severity groups: SC group exhibited a higher neutrophil fraction and a lower monocyte fraction than MM group, whereas lymphocyte, eosinophil and basophil proportions were indistinguishable (Figure S5). Around a week after admission, the distributions of all five cell types converged, and no between-group differences persisted (Figure S5), indicating that the leukocyte imbalance is a transitory feature from acute to recovery of the severe disease. After incorporating these cell-type covariates, 29 of the original 30 CpG-gene pairs lost statistical significance, demonstrating that most apparent methylation-expression associations were confounded by cell-type heterogeneity. The single CpG that retained significance—chr6:36697843—remained inversely associated with the expression of its neighbouring gene, *FKBP5* (Figure 2C, D; Table S4). The persistence of the chr6:36697843-*FKBP5 cis*-eQTM after rigorous adjustment establishes *FKBP5* as a cell-type proportion-independent epigenetic marker of disease severity.

### Resolution of acute-phase epigenetic alterations during COVID-19 recovery

We next examined whether the severity-associated epigenetic alterations observed during the acute phase were reversible during recovery. Specifically, we focused on the 30 CpGs and their corresponding 16 genes identified from the *cis*-eQTM analysis. Methylation and expression profiles were examined in the hospitalized MM and SC groups, an independent convalescent cohort (4–12 weeks post-infection, n = 90), and healthy controls (no viral infection at the time of collection, n = 344). A heatmap of the methylation *β*-values across all COVID-19 patients revealed a distinct group-wise separation during the acute phase, with SC patients displaying pronounced hyper- or hypo-methylation relative to MM group (Figure 3A). Notably, this severity-dependent stratification diminished substantially during the recovery phase, and methylation profiles became more similar between the SC and MM groups (Figure S6). To further assess the restoration trajectory, we visualized the dynamics using principal component analysis (PCA). In the gene expression PCA (Figure 3C), both SC and MM groups shifted almost completely toward the healthy cluster by 2–3 weeks post-infection, indicating rapid normalization of transcriptional responses (Figure S7). However, the DNA methylation PCA (Figure 3B) revealed that while MM group largely overlapped with convalescent and healthy controls, the SC group exhibited only a partial shift, especially along PC2. Although directional recovery was evident, the residual separation from the healthy baseline implies that full epigenetic reversion may require a longer time course, especially for those with severe disease.

**Figure 3.**
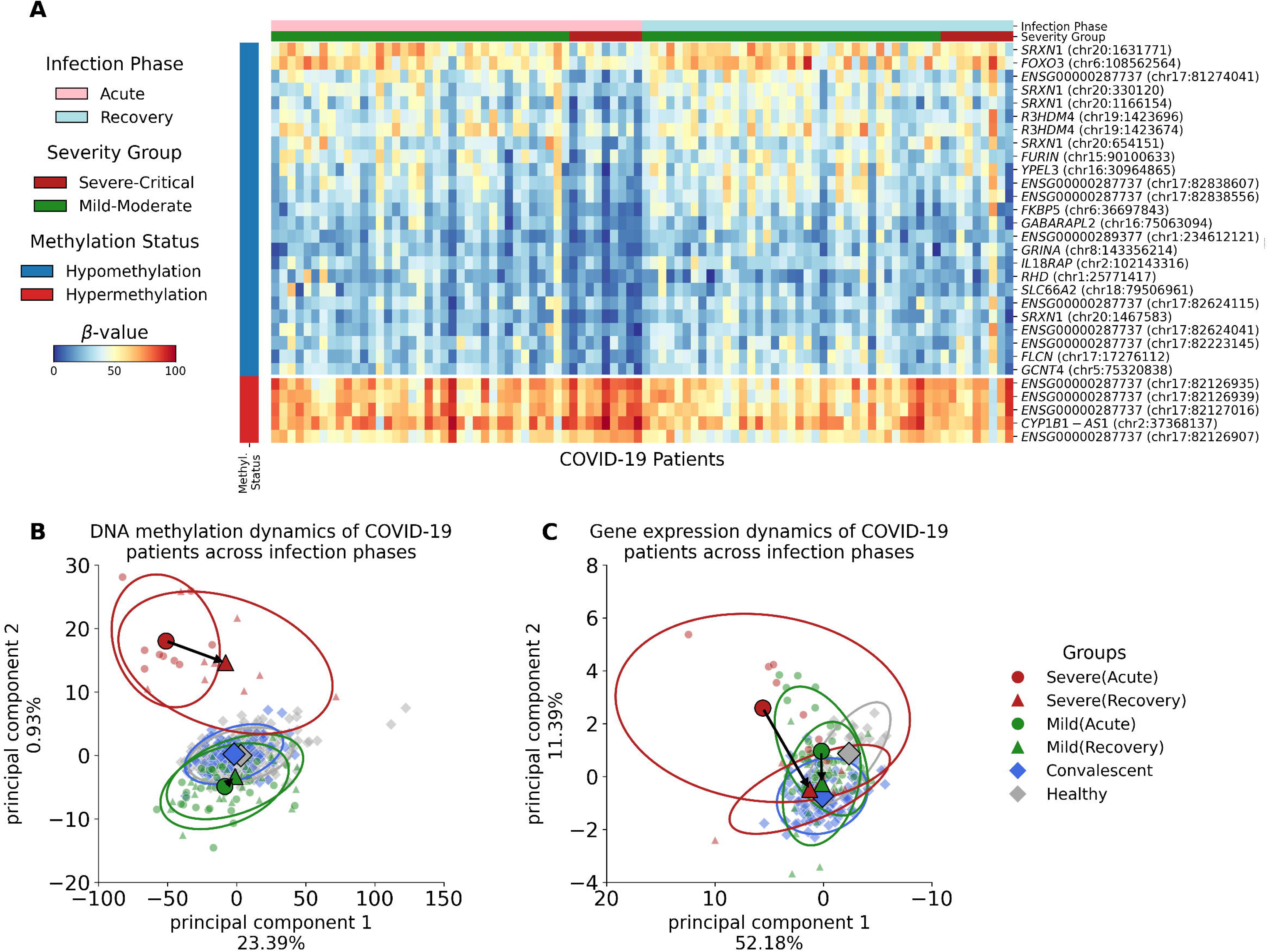
Multiomic Landscape of COVID-19 Recovery in Mild-Moderate and Severe-Critical Groups. **A)** Heatmap depicting DNA methylation profiles of *cis*-eQTM markers in COVID-19 patients across infection phases (acute vs. recovery) and severity groups (Mild-Moderate vs. Severe-Critical). Differentially methylated CpG sites are indicated by color: hypermethylated sites are shown in red and hypomethylated sites in blue. Infection phases are represented as acute (pink) and recovery (light blue), while severity groups are differentiated by color: Mild-Moderate (dark green) and Severe-Critical (dark red). The β-value scale, ranging from 0 to 100, indicates the proportion of methylation at each CpG site. **B)** Principal component analysis (PCA) of DNA methylation restoration between the acute and recovery phases in COVID-19 patients and healthy controls. **C)** PCA of gene expression restoration between acute and recovery phases in COVID-19 patients. Arrows track group-level transitions, with ellipses denoting 3 standard deviations from the group mean. Abbreviation: Methyl. Status, Methylation Status.

To examine individual-level changes, we analyzed the methylation difference between the acute and recovery phase in each SC group (Figure S8). Most patients exhibited significant *β-*value shifts across both hypermethylated and hypomethylated CpGs, consistent with the overall trend of epigenetic normalization. However, one patient (C19-C058) presented an exception. Unlike other patients in the SC group, this patient showed virtually no methylation change between two time points. Clinical metadata indicated that C19-C058 was classified as “Critical”, had a high Charlson Comorbidity Index (CCI = 5), and remained on oxygen therapy at the time of recovery-phase blood collection. The follow-up sample was collected just 10 days after baseline, which may have been insufficient time to capture epigenetic recovery.

## Discussion

Using early-phase whole-blood multi-omics from clinically well-characterized COVID-19 patients, we mapped CpG-to-transcript links with *cis*-eQTM analysis and uncovered a focused set of “reactive” CpGs whose methylation shifts reflected both disease severity and subsequent recovery. During the acute phase, coordinated hypo- or hyper-methylation at these sites corresponded to sharp transcriptional changes that distinguished severe-critical from mild-moderate cases. After correcting for transient leukocyte imbalances, only one CpG–RNA pair, *FKBP5,* remained independently associated with severity, highlighting it as a core epigenetic signal rather than a by-product of immune-cell redistribution. Longitudinal follow-up demonstrated that both methylation and gene expression at the reactive loci returned to baseline within 4–12 weeks, underscoring the dynamic and largely reversible nature of COVID-19-induced epigenetic programming. These findings establish a compact, cell-type composition–independent CpG/RNA signature that captures the trajectory from acute pathology to convalescence, providing possible prognostic biomarkers with mechanistic insight pertaining to host response and recovery during COVID-19.

By distinguishing between severe and mild cases based on oxygen therapy and intensive care unit (ICU) admission, we explored the molecular underpinnings associated with distinct clinical trajectories. Although there were only ten severe cases in this study, they represented 27% of the data, comparable to the proportion reported in previous large-scale studies. For example, a February 2020 study of 72,314 COVID-19 cases in China found that the majority (81%) of patients experienced mild to moderate symptoms, while 14% suffered from severe respiratory issues, and 5% progressed to critical conditions, including respiratory failure^15^.

This study is among the first to apply *cis*-eQTM analysis for dissecting gene regulatory programs linked to COVID-19 severity. We identified a distinct pattern of hypomethylation and concurrent upregulation of six genes—*SRXN1*, *FURIN*, *IL18RAP*, *FOXO3*, *GCNT4*, and *FKBP5*—involved in key processes, including host cell invasion, oxidative stress response, interleukin-18-mediated signaling, and lung fibrosis. These five genes are not only epigenetically regulated but also biologically implicated in known pathogenic mechanisms which are recognized as drivers of COVID-19 severity. Notably, *SRXN1*, which was hypomethylated and up-regulated in severe cases across five distinct CpG loci, encodes Sulfiredoxin 1, a critical antioxidant mitigating reactive oxygen species (ROS) damage. Its dysregulation amplifies oxidative stress and has been repeatedly reported to be highly expressed in both blood and lung tissues of severe COVID-19 patients^16–20^. *FURIN*, another gene up-regulated in severe cases, facilitates viral entry by cleaving the S1/S2 junction of the SARS-CoV-2 spike protein^21^. Its expression correlates with disease progression and has been identified as a potential target for antiviral therapy^22,23^. Our results also revealed significant upregulation of *IL18RAP* and *FOXO3* in severe COVID-19 cases—genes that are mechanistically linked to hyper-inflammatory responses and respiratory dysfunction. *IL18RAP*, a key mediator of IL-18 signaling, likely contributes to excessive immune activation and cardiopulmonary complications through inflammasome-driven IL-18 release, a pathway implicated in macrophage activation syndrome and multi-organ failure^24–26^. In parallel, *FOXO3*, which regulates immune balance and oxidative stress, was associated with increased oxygen demand and inflammatory lung injury, suggesting that its overexpression may exacerbate pulmonary damage in severe disease^27–29^.

Interestingly, *GCNT4* was the only gene found to be epigenetically downregulated in severe patients. A prior proteomic study reported its elevation in respiratory failure among severe COVID-19 cases^30^. This discrepancy may reflect differences in regulatory control between the transcriptome and proteome or patient cohort characteristics.

To account for the known shifts in immune cell proportions during acute infection, such as neutrophilia and lymphopenia, we repeated the *cis*-eQTM analysis while adjusting for estimated immune cell-type proportion. Most CpG–gene associations lost significance following this correction, suggesting that they were primarily driven by inflammation-induced cell redistribution. In contrast, the persistence of the *FKBP5* association after cell-type adjustment suggests that its regulation may reflect a cell-intrinsic epigenetic signal, strongly relevant to the pathophysiology of severe COVID-19. *FKBP5* encodes FKBP51, an Hsp90 co-chaperone that attenuates glucocorticoid-receptor (GR) signaling^31,32^. By modulating GR sensitivity it fine-tunes hypothalamic-pituitary-adrenal (HPA) feedback to stress and intersects with NF-κB/RIG-I pathways that shape innate-immune and inflammatory responses^33,34^. Transcriptomic studies show *FKBP5* up-regulation in corticosteroid-treated airway epithelium and in brains of fatal COVID-19 cases^35^, suggesting that GR-linked stress and treatment responses converge on *FKBP5* during advanced disease, although its causal role in COVID-19 severity remains inconclusive. While these findings point to a functional association between *FKBP5* activity and COVID-19 pathophysiology, further mechanistic and longitudinal studies are required to clarify whether *FKBP5* plays a driver, mediator, or consequence role in severe disease progression.

The remaining 10 *cis-*eQTM genes not covered in-depth in this study may still contribute to the broader molecular response to SARS-CoV-2 infection. Further investigation is warranted on elucidating their roles, which could reveal indirect effects on disease severity or related complications. Taken together, the blood-based multi-omic markers discovered here underscore pulmonary lesions and associated clinical symptoms. These markers could help predict respiratory complications and monitor disease severity.

In the recovery phase, spanning two to three weeks post-infection, we observed a relatively fast restoration of both the epigenome and transcriptome to levels akin to those of COVID-19 convalescence, across both severe and mild patient cohorts. The gene regulatory mechanisms observed in severe COVID-19 survivors, who avoided fatal outcomes, did not show significant disparities when compared to those in mild cases. This indicates that the identified CpGs are reactive to both the acute infection and recovery over the relatively short time period, reflecting the rapidly evolving pathological trajectory of the patients. Despite these overall trends, our findings also point to heterogeneity in recovery trajectories, particularly among severe-critical patients. In a subset of individuals, epigenetic reversal was less pronounced, and in one notable case (C19-C058), virtually no change in DNA methylation was observed between the acute and follow-up samples. This individual had a high Charlson Comorbidity Index (CCI), remained on oxygen therapy at follow-up, and their recovery-phase sample was taken just 10 days after the acute-phase time point. While these clinical factors may explain the apparent lack of epigenetic normalization, the case highlights that not all patients follow the same recovery timeline—and that molecular reversal may lag behind clinical resolution in more complex or prolonged disease courses.

In conclusion, these observations demonstrate that both cell-intrinsic and extrinsic epigenetic markers are responsive to COVID-19 pathophysiology and capable of tracking the acute-to-recovery transition. However, the variability across individuals also points to the importance of integrating clinical context and extending the observation window to fully understand the persistence, resolution, or relapse of molecular alterations—especially in relation to long-term sequelae, such as Long-COVID.

Our investigation encountered several critical limitations. First, the COVID-19 severe cases in this study exhibited variability in the timing and duration of oxygen therapy following hospitalization (Figure 1B). This discrepancy poses constraints on elucidating the exact association between identified biomarkers and the severity of the condition. Second, the limited number of samples in the severe group presents a challenge in drawing comprehensive general conclusions. Third, the lack of standardized control over the recovery phase, ranging from two to three weeks per patient, introduces variability that influences the interpretation of results. Additionally, the separate sample collection periods in 2021 and 2022 renders complexity. Different SARS-CoV-2 sub-strains – Delta and Omicron – were predominant during these times. The variants differed in transmissibility, virulence, and immune evasion, likely influencing disease severity and treatment response^36^. Lastly, while cytokine measurements were not performed in this study, the absence of these data limits our ability to directly evaluate the contribution of cytokine-mediated inflammatory responses to disease severity. Given the established role of cytokine storms in severe COVID-19 cases, future studies integrating cytokine profiling with epigenetic and transcriptomic data would offer a more comprehensive understanding of immune dysregulation in acute infection.

## Materials and methods

### Samples and clinical characteristics

We collected whole blood samples from 46 patients diagnosed with COVID-19 from Ulsan University Hospital (UUH), Ulsan, Republic of Korea. This study has received approval from the Institutional Review Board (IRB) of UUH and Ulsan National Institute of Science and Technology (UNIST) (IRB No.: UUH-2021-04-011-004, UNISTIRB-21-15-A).

For the hospitalized patient group, whole blood samples were collected twice: once during the acute phase (at the time of hospital admission) and once during the recovery phase (2 to 3 weeks after hospitalization). For the convalescent patient group, whole blood samples were obtained between 4 to 12 weeks after COVID-19 diagnosis. These individuals had been previously hospitalized due to COVID-19 but had fully recovered at the time of sample collection. Our sampling period is divided into two halves. The first was during 2021 which was followed by the second in 2022. The acute phase of infection was defined as the period from hospitalization due to COVID-19 to one week thereafter, while the subsequent period of up to three weeks was defined as the recovery phase. A healthy control group consisted of 344 whole blood samples collected (309 samples for methylation, 34 for gene expression studies, and one sample for both omics) from the Korean Genome Project (KGP), approved by the IRB at UNIST in Ulsan, South Korea (IRB No.: UNISTIRB-21-66-A).

### Patient classification of COVID-19 severity

The patients were assigned their severity upon diagnosis according to Food and Drug Administration (FDA) severity categorization and categorized into four groups: “Mild”, “Moderate”, “Severe”, and “Critical” (Table S5) ^37,38^. We further categorized the patients upon analysis defining their severity into two discrete groups: “Mild-Moderate” (MM) for mild and moderate categories and “Severe-Critical” (SC) for severe and critical categories. Severe-critical patients suffered from respiratory symptoms with 80% of the patients requiring oxygen therapies, such as nasal prong or high-flow nasal cannula, or both, as compared to 14.6% for the MM group. In addition, 40% of the SC cases were admitted to the ICU, while none in the MM (Table 2).

**Table 2.** Baseline Characteristics of COVID-19 Hospitalized Patients, Convalescent Patients, and Non-infected Healthy Individuals. Demographic and clinical data are summarized for the clinical cohorts. Continuous variables (e.g., age and BMI) are presented as mean values with minimum and maximum ranges in parentheses. Categorical variables (e.g., sex, smoking status, and CCI) are expressed as counts with corresponding percentages. Smoking status was self-reported and categorized as current, former, or never smoker. Abbreviations: HFNC, high-flow nasal cannula; ARDS, acute respiratory distress syndrome; CCI, Charlson Comorbidity Index.

|  | COVID-19 Hospitalized patients |  | Convalescent group (n=90) | Healthy controls (n=310) |
| --- | --- | --- | --- | --- |
|  | Mild-Moderate group (n=37) | Severe-Critical group (n=9) |  |  |
| Age, mean(min, max) | 44(19,71) | 50(24,66) | 42(19,66) | 42(19, 66) |
| Sex , n(%) |  |  |  |  |
| Male | 24(64.9) | 4(44.4) | 45(50) | 188(60) |
| Female | 13(35.1) | 5(55.6) | 45(50) | 122(40) |
| **BMI** | 24.8 | 25.5 | 24.4 | 23.45 |
| **Smoking status, n(%)** |  |  |  |  |
| Current | 3(8.1) | 3(33.3) |  | 46(14.8) |
| Former | 3(8.1) | 1(11.1) |  | 79(25.5) |
| Never | 31(83.8) | 5(55.6) |  | 173(55.8) |
| NA | 0 | 0 | 90 | 12(3.9) |
| **Oxygen therapy, n(%)** |  |  |  |  |
| Yes |  |  |  |  |
| Nasal prong only | 8(21.6) | 5(55.6) |  |  |
| HFNC only | 0 | 2(22.2) |  |  |
| Both (Nasal prong and HFNC) | 0 | 2(22.2) |  |  |
| No | 26(70.3) | 0 |  |  |
| **Intensive care unit, n(%)** |  |  |  |  |
| Yes | 0 (0) | 3(33.3) |  |  |
| No | 37(100) | 6(66.7) |  |  |
| **Use of antiviral drugs, n(%)** |  |  |  |  |
| Yes |  |  |  |  |
| Regdanvimab | 3(8.1) | 2(22.2) |  |  |
| Remdesivir | 10(27) | 6(66.7) |  |  |
| No | 24(64.9) | 1(11.1) |  |  |
| **Use of steroid, n(%)** |  |  |  |  |
| Yes | 11(29.7) | 7(77.8) |  |  |
| No | 26(70.3) | 2(22.2) |  |  |
| **Pneumonia, n(%)** |  |  |  |  |
| Yes | 10 (27) | 7(77.8) |  |  |
| No | 27 (73.0) | 2(22.2) |  |  |
| **ARDS, n(%)** |  |  |  |  |
| Yes | 0 (0) | 2(22.2) |  |  |
| No | 37(100) | 7(77.8) |  |  |
| **Vaccination status, n(%)** |  |  |  |  |
| Yes | 13(35.1) | 6(66.7) |  |  |
| No | 24(64.9) | 3(33.3) |  |  |
| **CCI, n(%)** |  |  |  |  |
| 0-1 | 24(64.9) | 1(11.1) |  |  |
| 2 | 4(10.8) | 4(44.4) |  |  |
| ≥ 3 | 8(21.6) | 3(33.3) |  |  |

### Clinical Information

We collected clinical data from informed and consenting patients alongside whole blood samples. The dataset included routine laboratory measurements, smoking history, and viral load estimates based on PCR cycle threshold (Ct) values (Figure S1). To investigate clinical correlates of disease severity, we compared these parameters across patient groups stratified by COVID-19 severity. Clinical variables with insufficient sample size (fewer than three observations in any group) were excluded from downstream analyses to ensure statistical robustness.

### Target Bisulfite Sequencing and Methylation Quantification

Frozen whole blood samples collected in EDTA tubes (Becton Dickinson, #367856) were used for genomic DNA (gDNA) extraction, performed using the DNeasy Blood & Tissue Kit (Qiagen, #69506) according to the manufacturer’s instructions.

Bisulfite-converted DNA libraries were prepared using the SureSelectXT Methyl-Seq Target Enrichment System Kit (Agilent, #G9651), following the manufacturer’s protocol. Sequencing was performed on the NovaSeq 6000 platform (Illumina), generating 150 bp paired-end reads. Raw sequencing reads were processed using Fastp (v0.23.1) to remove adapters and low-quality bases^39^, with the following parameters: --cut_front 1 --cut_right 0 --cut_tail 1 -- detect_adapter_for_pe --trim_front1 15 --trim_front2 15 --trim_tail1 0 --trim_tail2 0 -- cut_mean_quality 20 --n_base_limit 1 --average_qual 20 34. Read quality was assessed before and after filtering reads using FastQC (v0.11.9)^40^.

Filtered reads were aligned to the human reference genome (GRCh38.p13) using Bismark (v0.23.1), a bisulfite-aware aligner^41^. Methylation calling and quantification were conducted using the methylKit R package (v1.20.0)^42^. To annotate CpG sites and associate them with proximal genes or regulatory features, we used the annotatr R package (v1.20.0)^43^.

### mRNA Sequencing and Expression Quantification

Frozen whole blood samples collected in PAXgene® Blood RNA Tubes (PreAnalytiX, #762174) were used for total RNA extraction. RNA integrity and concentration were assessed using a Qubit 2.0 Fluorometer (Thermo Fisher Scientific, #Q32866) and the Qubit RNA HS Assay Kit (Thermo Fisher Scientific, #Q32854). For mRNA enrichment, 200 ng of total RNA per sample was processed using the Dynabeads mRNA Purification Kit (Thermo Fisher Scientific, #01152851), which depletes rRNA and isolates polyadenylated transcripts using oligo(dT) beads.

Library preparation was performed using the MGIEasy RNA Directional Library Prep Set (MGI, #MG1000006386), following the manufacturer’s protocol. Fragment size distribution was checked using 4150 TapeStation (Agilent, #G2992A) with the cDNA D1000 ScreenTape (Agilent, #5067-5582). Final libraries were quantitated using the Qubit 2.0 Fluorometer (Thermo Fisher Scientific, Q32866). RNA sequencing was run on DNBSEQ-T7RS (MGI) platform, generating 150 bp paired-end reads.

RNA-seq reads were trimmed and quality-filtered using Fastp (v0.23.1) with the same parameters as for DNA methylation processing. Read quality was verified using FastQC (v0.11.9).

Filtered reads were aligned to the human reference genome (GRCh38.p13) using STAR (v2.7.10b), a splice-aware aligner, with default settings^44^. Gene-level quantification was performed using RSEM (v1.3.3) using default options^45^, with gene annotations from GENCODE v42 (GFF3 format). Raw expression values were derived from the expected counts generated by RSEM.

### Batch Effect Correction of Gene Expression Data

To integrate gene expression data from an independent cohort of 35 healthy individuals, we implemented rigorous batch effect correction to ensure comparability. All samples were processed using an identical bioinformatic pipeline as RNA-seq data for COVID-19 patients to avoid potential *in silico* batch effects. We applied the ComBat-seq method implemented in the sva R package (v3.54.0)^46^ to remove batch effect by differences in sequencing platforms (i.e., NovaSeq 6000 vs DNBSEQ-T7RS) while preserving the biological condition (i.e., Healthy vs. COVID-19 status).

### Normalization of Gene Expression Data

To account for differences in sequencing depth and library composition, raw expected gene counts were normalized using the DESeq2 R package (v1.38.3)^47^. A variance stabilizing transformation (VST) was subsequently applied to the normalized counts. VST approximates a log2 transformation while decoupling mean-variance relationships of RNA-seq data, thereby improving the accuracy and interpretability of downstream statistical analyses. VST-transformed values were used for pairwise comparisons stratified by COVID-19 severity, allowing expression differences to be interpreted in fold-change-like units. These transformed values were also used for principal component analysis (PCA) to assess global expression patterns and sample clustering.

### Discovery of Differential Methylation CpGs (DMC) Associated with COVID-19 Severity

Methylkit (R package; v1.20.0)^42^ was used to discover the Differentially Methylated CpGs (DMC) for COVID-19 severity, treating age and sex of the patients as covariates. We utilized the LOO (leave-one-out) method for cross-validation, iteratively collecting DMC without a sample for each round of discovery. Methylation sites that suffice the thresholds of absolute methylation difference (|meth.diff|) > 10 and FDR < 0.05 were selected as significant markers of the severity. The adjustment for *P*-value was done by Benjamini-Hochberg correction. We only selected a set of markers, those of which overlapped over all the folds.

### Leukocyte-adjusted Discovery of DMCs Associated with COVID-19 Severity

We repeated the leave-one-out (LOO)-based DMC discovery as previously described. Here, we adjusted for age, sex, and blood cell-type composition (%)—including neutrophils, lymphocytes, monocytes, basophils, and eosinophils. This analysis included only samples with available cell count measurements recorded during clinical data collection, comprising 24 MM and 9 SC patients. The significance thresholds and marker selection criteria were identical to those used in the leukocyte-unadjusted analysis.

### Discovery of Differentially Expressed Genes (DEGs) Associated with COVID-19 Severity

Differentially Expressed Genes (DEGs) associated with COVID-19 severity were identified using the DESeq2 R package (v1.38.3). Given the limited number of Severe-Critical (SC) cases (n=9), we employed a leave-one-out (LOO) cross-validation strategy to enhance the robustness of DEG discovery. In each iteration, one SC sample was excluded, and differential expression analysis was performed between the remaining SC and mild cases. This procedure was repeated nine times, each time omitting a different SC sample. Genes were considered significantly differentially expressed if they satisfied the following thresholds: absolute log₂ fold change (|log₂FoldChange|) > 1.3 and false discovery rate (FDR) < 0.05. To define robust DEGs (DEG-LOOs), we retained only those genes that exhibited consistent directionality—either upregulation or downregulation in SC patients—in at least seven out of the nine independent LOO iterations.

### Gene Ontology Enrichment of DMCs and DEGs

To interpret the biological functions associated with differential DNA methylation and gene expression associated with COVID-19 severity, Gene Ontology (GO) enrichment analysis was conducted separately for genes linked to differentially methylated CpGs (DMCs) and differentially expressed genes (DEGs). GO enrichment analysis was performed using ShinyGO v0.82 (https://bioinformatics.sdstate.edu/go82/)^48^, focusing on the Biological Process (GO:BP) category. The gene universe was defined as all genes expressed in the RNA-seq dataset (for DEGs) or all genes targeted by measured CpGs (for DMCs). Statistical significance was assessed using a hypergeometric test with Benjamini–Hochberg correction. GO terms with FDR < 0.05 were considered significantly enriched.

### Prioritization of Severity-Associated CpG Sites via *cis*-eQTM (Expression Quantitative Trait Methylation) Analysis

We performed an Expression Quantitative Trait Methylation (eQTM) analysis to assess the relationship between CpG methylation and gene expression (DESeq2-normalized read count) that have been discovered during previous steps (i.e., DMC and DEG) (Figure S9). Spearman correlation was used to quantify this relationship, with a significance threshold of absolute correlation coefficient > 0.5 using scipy.stats (v1.4.1). Gene coordinates and transcription start sites (TSS) were extracted from GENCODE v42 (GTF format). For *cis*-association analysis, we restricted the search space to CpG sites located within a ±1Mbp window of the corresponding gene’s TSS. This window was selected to capture local regulatory effects of CpG methylation on gene expression.

### Statistical analysis and Visualization

Welch’s *t*-test (a two-sample independent t-test of unequal variances) was performed to find the mean difference of methylation values between different groups of infection stages using scipy.stats (v1.4.1). The normality assumption for the distributions of omics data was checked by the Shapiro-Wilk test using scipy.stats (v1.4.1) (Table S6, 7). The *P*-values were adjusted with the Benjamini-Hochberg approach using statsmodels.stats (v0.13.2). Wilcoxon rank-sum test using scipy.stats (v1.4.1) was performed to test the statistically significant differences in gene expression and clinical values across severities. Principal components for DNA methylation and gene expression were computed using sklearn.decomposition.PCA (v0.23.2). All visualizations were drawn by using matplotlib (v3.5.3) and seaborn (v0.11.0).

## Data Availability

Raw sequencing data and materials used in the study are available from the corresponding author upon request.

## Code Availability

The codes used to generate data and calculate statistics are openly available in the Github page: https://github.com/korean-genomics-center/Multiomics_COVID19_Severity

## Author contributions

Hyojung Ryu, Conceptualization, Project administration, Formal analysis, Data curation, Investigation, Methodology, Results interpretation, Visualization, Writing – original draft, Writing – review and editing; Kyungwhan An, Formal analysis, Data curation, Investigation, Methodology, Results interpretation, Visualization, Writing – original draft, Writing – review and editing; Yoonsung Kwon, Data curation, Formal analysis, Methodology, Writing – review and editing; Yeonsu Jeon, Writing – review and editing; Sungwon Jeon, Writing – review and editing; Hansol Choi, Conceptualization, Project administration; Yeo Jin Kim, Writing – review and editing; Sunhwa Kim, Patient recruitment and sample collection, Clinical phenotyping, Data curation; Ok Joo Sul, Sample processing, Data curation; SangJoon Lee, Writing – review and editing; Asaph Young Chun, Writing – review and editing; Eun-Seok Shin, Conceptualization, Results interpretation, Supervision, Writing – original draft, Writing – review and editing; Seung Won Ra, Conceptualization, Patient recruitment and sample collection, Data curation, Project administration, Conceived and designed the study, Curated clinical phenotyping data, Writing – review and editing; Jong Bhak, Conceptualization, Results interpretation, Supervision, Conceived and designed the study, Writing – review and editing, Funding acquisition

## Acknowledgements

We thank all voluntary participants for donating their blood and the city of Ulsan for supporting the project. We thank Asta Blazyte for her assistance in proofreading and refining the English of this manuscript. We also appreciate the Ulsan ICT Promotion Agency (UIPA), which provided us with the BioDataFarm system, which supports the storage, analysis, and management of the BioBigData.

## References

1. Mohandas, S., et al. Immune mechanisms underlying COVID-19 pathology and post-acute sequelae of SARS-CoV-2 infection (PASC). Elife 12(2023).

2. Karki, R., et al. ZBP1-dependent inflammatory cell death, PANoptosis, and cytokine storm disrupt IFN therapeutic efficacy during coronavirus infection. Sci Immunol 7, eabo6294 (2022).

3. Montazersaheb, S., et al. COVID-19 infection: an overview on cytokine storm and related interventions. Virol J 19, 92 (2022).

4. Barturen, G., et al. Whole blood DNA methylation analysis reveals respiratory environmental traits involved in COVID-19 severity following SARS-CoV-2 infection. Nat Commun 13, 4597 (2022).

5. Qin, W., Scicluna, B.P. & van der Poll, T. The Role of Host Cell DNA Methylation in the Immune Response to Bacterial Infection. Front Immunol 12, 696280 (2021).

6. Castro de Moura, M., et al. Epigenome-wide association study of COVID-19 severity with respiratory failure. EBioMedicine 66, 103339 (2021).

7. Bradic, M., et al. DNA methylation predicts the outcome of COVID-19 patients with acute respiratory distress syndrome. J Transl Med 20, 526 (2022).

8. Li, Y.Y., et al. Cell-free DNA methylation reveals cell-specific tissue injury and correlates with disease severity and patient outcomes in COVID-19. Clin Epigenetics 16, 37 (2024).

9. Balnis, J., et al. Blood DNA methylation and COVID-19 outcomes. Clin Epigenetics 13, 118 (2021).

10. Pacis, A., et al. Bacterial infection remodels the DNA methylation landscape of human dendritic cells. Genome Res 25, 1801–1811 (2015).

11. Salgado-Albarran, M., et al. Comparative transcriptome analysis reveals key epigenetic targets in SARS-CoV-2 infection. NPJ Syst Biol Appl 7, 21 (2021).

12. Keshawarz, A., et al. Expression quantitative trait methylation analysis elucidates gene regulatory effects of DNA methylation: the Framingham Heart Study. Sci Rep 13, 12952 (2023).

13. Kim, S., et al. Cis- and trans-eQTM analysis reveals novel epigenetic and transcriptomic immune markers of atopic asthma in airway epithelium. J Allergy Clin Immunol 152, 887–898 (2023).

14. Ruiz-Arenas, C., et al. Identification of autosomal cis expression quantitative trait methylation (cis eQTMs) in children’s blood. Elife 11(2022).

15. Wu, Z. & McGoogan, J.M. Characteristics of and Important Lessons From the Coronavirus Disease 2019 (COVID-19) Outbreak in China: Summary of a Report of 72 314 Cases From the Chinese Center for Disease Control and Prevention. JAMA 323, 1239–1242 (2020).

16. Saheb Sharif-Askari, N., et al. Upregulation of oxidative stress gene markers during SARS-COV-2 viral infection. Free Radic Biol Med 172, 688–698 (2021).

17. Xia, J., et al. Immune Response Is Key to Genetic Mechanisms of SARS-CoV-2 Infection With Psychiatric Disorders Based on Differential Gene Expression Pattern Analysis. Front Immunol 13, 798538 (2022).

18. Jain, S.K., et al. The potential link between inherited G6PD deficiency, oxidative stress, and vitamin D deficiency and the racial inequities in mortality associated with COVID-19. Free Radic Biol Med 161, 84–91 (2020).

19. Bastin, A., et al. Severity of oxidative stress as a hallmark in COVID-19 patients. Eur J Med Res 28, 558 (2023).

20. Gain, C., Song, S., Angtuaco, T., Satta, S. & Kelesidis, T. The role of oxidative stress in the pathogenesis of infections with coronaviruses. Front Microbiol 13, 1111930 (2022).

21. Peacock, T.P., et al. The furin cleavage site in the SARS-CoV-2 spike protein is required for transmission in ferrets. Nat Microbiol 6, 899–909 (2021).

22. Essalmani, R., et al. Distinctive Roles of Furin and TMPRSS2 in SARS-CoV-2 Infectivity. J Virol 96, e0012822 (2022).

23. Hossain, M.S., et al. Prediction of the Effects of Variants and Differential Expression of Key Host Genes ACE2, TMPRSS2, and FURIN in SARS-CoV-2 Pathogenesis: An In Silico Approach. Bioinform Biol Insights 15, 11779322211054684 (2021).

24. Liang, S., et al. SARS-CoV-2 spike protein induces IL-18-mediated cardiopulmonary inflammation via reduced mitophagy. Signal Transduct Target Ther 8, 108 (2023).

25. Nasser, S.M.T., et al. Elevated free interleukin-18 associated with severity and mortality in prospective cohort study of 206 hospitalised COVID-19 patients. Intensive Care Med Exp 11, 9 (2023).

26. Korotaeva, A.A., et al. Ratios between the Levels of IL-18, Free IL-18, and IL-1beta-Binding Protein Depending on the Severity and Outcome of COVID-19. Bull Exp Biol Med 176, 423–427 (2024).

27. Cheema, P.S., Nandi, D. & Nag, A. Exploring the therapeutic potential of forkhead box O for outfoxing COVID-19. Open Biol 11, 210069 (2021).

28. Gimenes-Junior, J., et al. FOXO3a regulates rhinovirus-induced innate immune responses in airway epithelial cells. Sci Rep 9, 18180 (2019).

29. Sullivan, J.A., Kim, E.H., Plisch, E.H. & Suresh, M. FOXO3 regulates the CD8 T cell response to a chronic viral infection. J Virol 86, 9025–9034 (2012).

30. Palmos, A.B., et al. Proteome-wide Mendelian randomization identifies causal links between blood proteins and severe COVID-19. PLoS Genet 18, e1010042 (2022).

31. Binder, E.B. The role of FKBP5, a co-chaperone of the glucocorticoid receptor in the pathogenesis and therapy of affective and anxiety disorders. Psychoneuroendocrinology 34 **Suppl 1**, S186–195 (2009).

32. Wochnik, G.M., et al. FK506-binding proteins 51 and 52 differentially regulate dynein interaction and nuclear translocation of the glucocorticoid receptor in mammalian cells. J Biol Chem 280, 4609–4616 (2005).

33. Hao, W., Wang, L. & Li, S. FKBP5 Regulates RIG-I-Mediated NF-kappaB Activation and Influenza A Virus Infection. Viruses 12(2020).

34. Erlejman, A.G., et al. NF-kappaB transcriptional activity is modulated by FK506-binding proteins FKBP51 and FKBP52: a role for peptidyl-prolyl isomerase activity. J Biol Chem 289, 26263–26276 (2014).

35. Green, R., et al. SARS-CoV-2 infection increases the gene expression profile for Alzheimer’s disease risk. Mol Ther Methods Clin Dev 27, 217–229 (2022).

36. Carabelli, A.M., et al. SARS-CoV-2 variant biology: immune escape, transmission and fitness. Nat Rev Microbiol 21, 162–177 (2023).

37. COVID-19: Developing Drugs and Biological Products for Treatment or Prevention. (Center for Drug Evaluation and Research, U.S. Food and Drug Administration, 2023).

38. Toussi, S.S., Hammond, J.L., Gerstenberger, B.S. & Anderson, A.S. Therapeutics for COVID-19. Nat Microbiol 8, 771–786 (2023).

39. Chen, S., Zhou, Y., Chen, Y. & Gu, J. fastp: an ultra-fast all-in-one FASTQ preprocessor. Bioinformatics 34, i884–i890 (2018).

40. Andrews, S. FastQC: a quality control tool for high throughput sequence data. (Cambridge, United Kingdom, 2010).

41. Krueger, F. & Andrews, S.R. Bismark: a flexible aligner and methylation caller for Bisulfite-Seq applications. Bioinformatics 27, 1571–1572 (2011).

42. Akalin, A., et al. methylKit: a comprehensive R package for the analysis of genome-wide DNA methylation profiles. Genome Biol 13, R87 (2012).

43. Cavalcante, R.G. & Sartor, M.A. annotatr: genomic regions in context. Bioinformatics 33, 2381–2383 (2017).

44. Dobin, A., et al. STAR: ultrafast universal RNA-seq aligner. Bioinformatics 29, 15–21 (2013).

45. Li, B. & Dewey, C.N. RSEM: accurate transcript quantification from RNA-Seq data with or without a reference genome. BMC Bioinformatics 12, 323 (2011).

46. Zhang, Y., Parmigiani, G. & Johnson, W.E. ComBat-seq: batch effect adjustment for RNA-seq count data. NAR Genom Bioinform 2, lqaa078 (2020).

47. Love, M.I., Huber, W. & Anders, S. Moderated estimation of fold change and dispersion for RNA-seq data with DESeq2. Genome Biol 15, 550 (2014).

48. Ge, S.X., Jung, D. & Yao, R. ShinyGO: a graphical gene-set enrichment tool for animals and plants. Bioinformatics 36, 2628–2629 (2020).

