## Supplemental materials for "Identification of reactive CpGs and RNA expression in early COVID-19 through *cis*-eQTM analysis reflecting disease severity and recovery"

### Supplementary Information

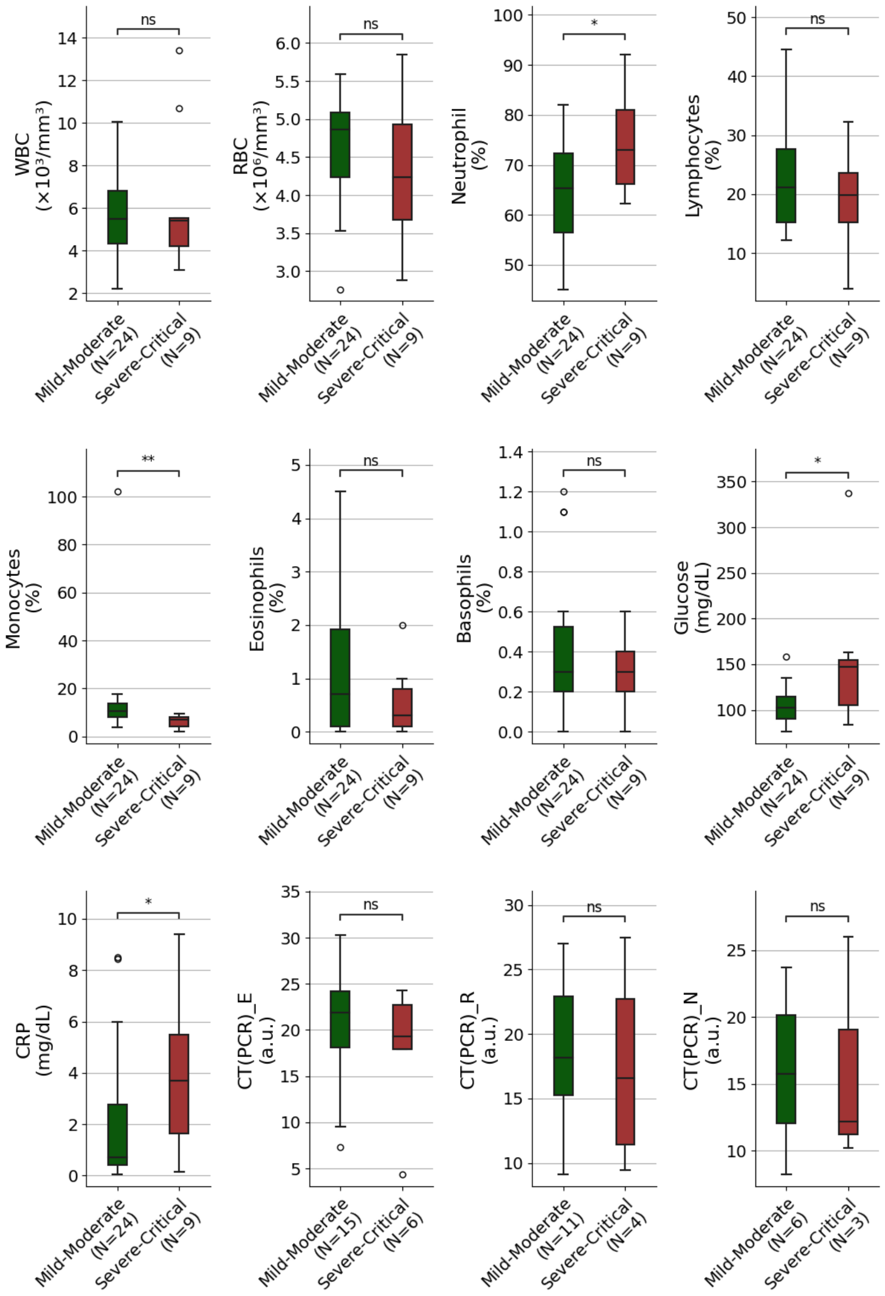

**Supplementary Fig 1. Clinical Characteristics of Mild–Moderate and Severe–Critical COVID-19 Patients During the Acute Phase.** The x-axis denotes severity groups, with sample sizes indicated in parentheses. The y-axis indicates the clinical parameter measured, with units denoted in parentheses. Boxplots display the median (centre line), interquartile range or IQR (box limits), and 1.5×IQR (whiskers), and outliers (open circles). Significance thresholds are indicated as follows: ns ($P \geq 0.05$), * ($P < 0.05$), ** ($P < 0.01$), *** ($P < 0.001$), **** ($P < 0.0001$). Abbreviations: WBC, White Blood Cell count; RBC, Red Blood Cell count; CRP, C-reactive protein concentration; CT(PCR), Cycle threshold (Ct) values from PCR assays for SARS-CoV-2 targets (*E*, *R*, and *N* genes); a.u., arbitrary units.

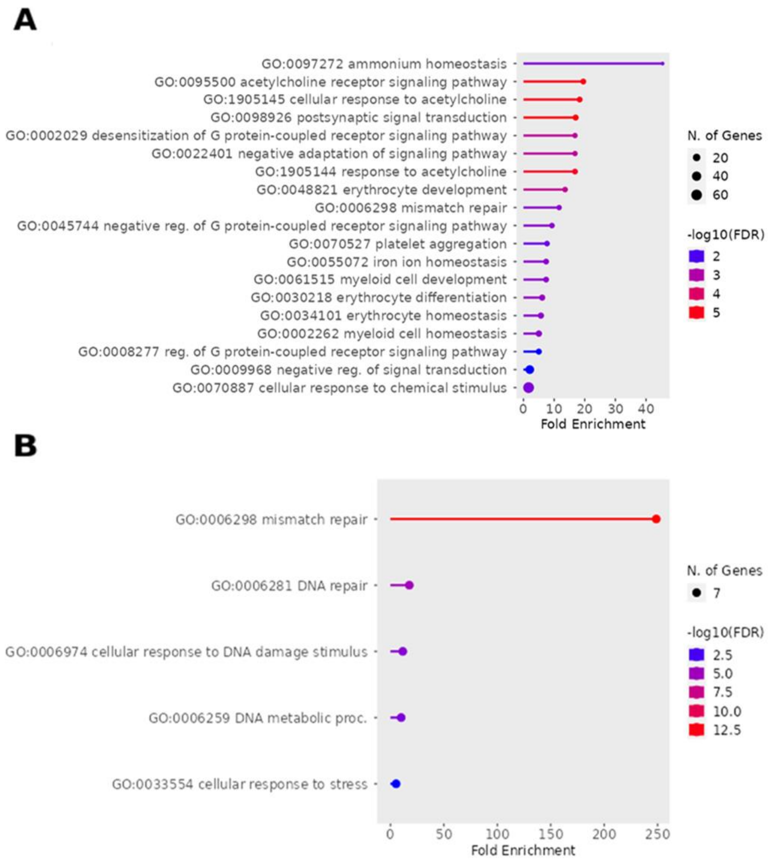


**Supplementary Fig 2. Gene Ontology Enrichment of 297 Severity-Associated DEG-LOOs During Acute COVID-19.** Lollipop plot illustrating Gene Ontology-Biological Process (GO-BP) enrichment for **A)** 283 up-regulated and **B)** 14 down-regulated genes linked to COVID-19 severity during the acute phase. The x-axis denotes fold enrichment, while the y-axis lists enriched GO terms. Dot size corresponds to the number of genes in each GO term, and color indicates statistical significance as –log10(FDR). Enrichment was performed using ShinyGO v0.82 based on the hypergeometric test with Benjamini–Hochberg correction, using all RNA-seq expressed genes as the background gene universe. Abbreviations: GO, Gene Ontology; BP, Biological Process; FDR, False Discovery Rate.

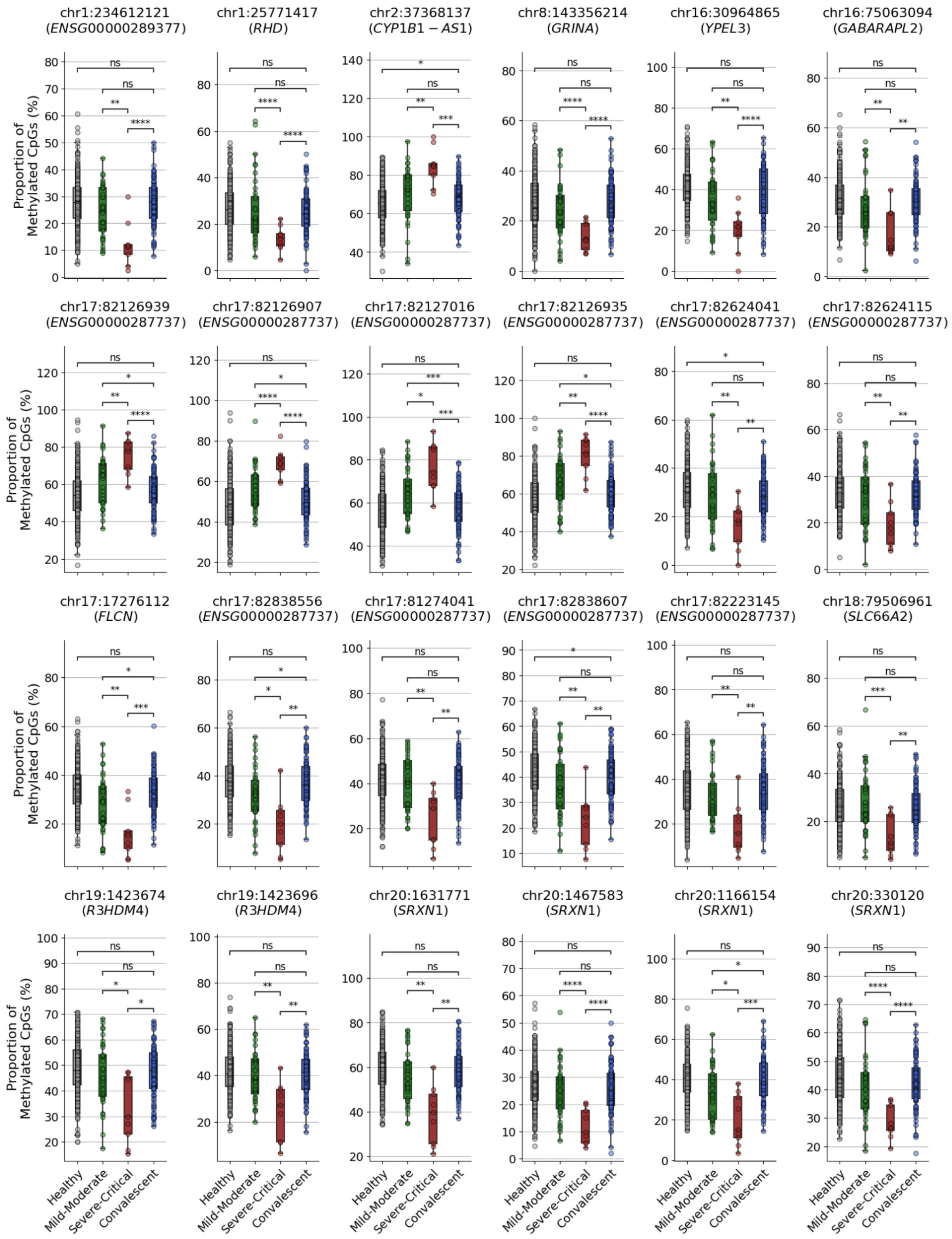
chr1:234612121 (ENSG00000289377)
chr1:25771417 (RHD)
chr2:37368137 (CYP1B1 − AS1)
chr8:143356214 (GRINA)
chr16:30964865 (YPEL3)
chr16:75063094 (GABARAPL2)
chr17:82126939 (ENSG00000287737)
chr17:82126907 (ENSG00000287737)
chr17:82127016 (ENSG00000287737)
chr17:82126935 (ENSG00000287737)
chr17:82624041 (ENSG00000287737)
chr17:82624115 (ENSG00000287737)
chr17:17276112 (FLCN)
chr17:82838556 (ENSG00000287737)
chr17:81274041 (ENSG00000287737)
chr17:82838607 (ENSG00000287737)
chr17:82223145 (ENSG00000287737)
chr18:79506961 (SLC66A2)
chr19:1423674 (R3HDM4)
chr19:1423696 (R3HDM4)
chr20:1631771 (SRXN1)
chr20:1467583 (SRXN1)
chr20:1166154 (SRXN1)
chr20:330120 (SRXN1)
Proportion of Methylated CpGs (%)
Healthy
Mild-Moderate
Severe-Critical
Convalescent

**Supplementary Fig 3. DNA Methylation Levels of 24 *cis*-eQTM CpGs Associated with COVID-19 Severity During the Acute Phase.** DNA methylation levels (% methylated CpGs) are shown for 24 *cis*-eQTM CpGs across four clinical groups: Healthy controls (n = 310), patients with Mild-Moderate (n = 37) or Severe-Critical (n = 9) COVID-19 during the acute phase, and Convalescent individuals (n = 90). The x-axis denotes severity groups. The y-axis represents the proportion of methylation at each CpG site. Boxplots display the median (centre line), IQR (box limits), 1.5×IQR (whiskers), and individual data points (closed circles). Statistical comparisons were conducted using Welch's *t*-test. Statistical significance is indicated as follows: ns ($P \geq 0.05$), * ($P < 0.05$), ** ($P < 0.01$), *** ($P < 0.001$), **** ($P < 0.0001$).

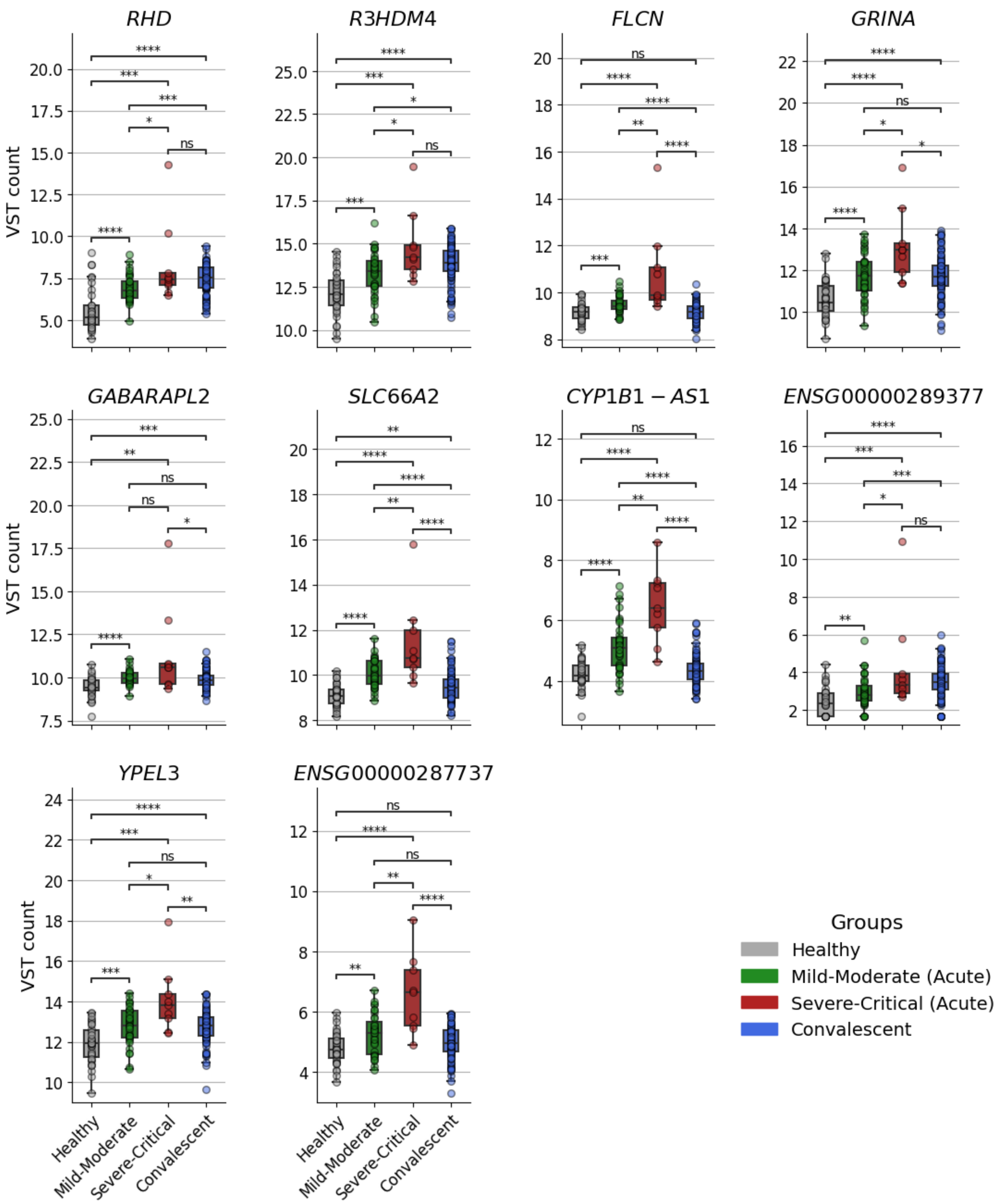
RHD
R3HDM4
FLCN
GRINA
GABARAPL2
SLC66A2
CYP1B1 − AS1
ENSG00000289377
YPEL3
ENSG00000287737
VST count
Healthy
Mild-Moderate
Severe-Critical
Convalescent
Groups
Healthy
Mild-Moderate (Acute)
Severe-Critical (Acute)
Convalescent

**Supplementary Fig 4. Expression Levels of 10 *cis*-eQTM-Regulated Genes Associated with COVID-19 Severity During the Acute Phase.** Expression levels of 10 genes regulated by *cis*-eQTM methylation sites are shown across four clinical groups: Healthy controls (n = 35), patients with Mild-Moderate (n = 36) or Severe-Critical (n = 9) COVID-19 during the acute phase, and Convalescent individuals (n = 90). The x-axis represents clinical groups.The y-axis shows gene expression values, represented as variance-stabilizing transformed (VST) counts. Boxplots display the median (centre line), IQR (box limits), 1.5×IQR (whiskers), and individual data points (closed circles). Statistical comparisons were conducted using the Wilcoxon rank-sum test. Statistical significance is denoted as follows: ns ($P \geq 0.05$), * ($P < 0.05$), ** ($P < 0.01$), *** ($P < 0.001$), **** ($P < 0.0001$).

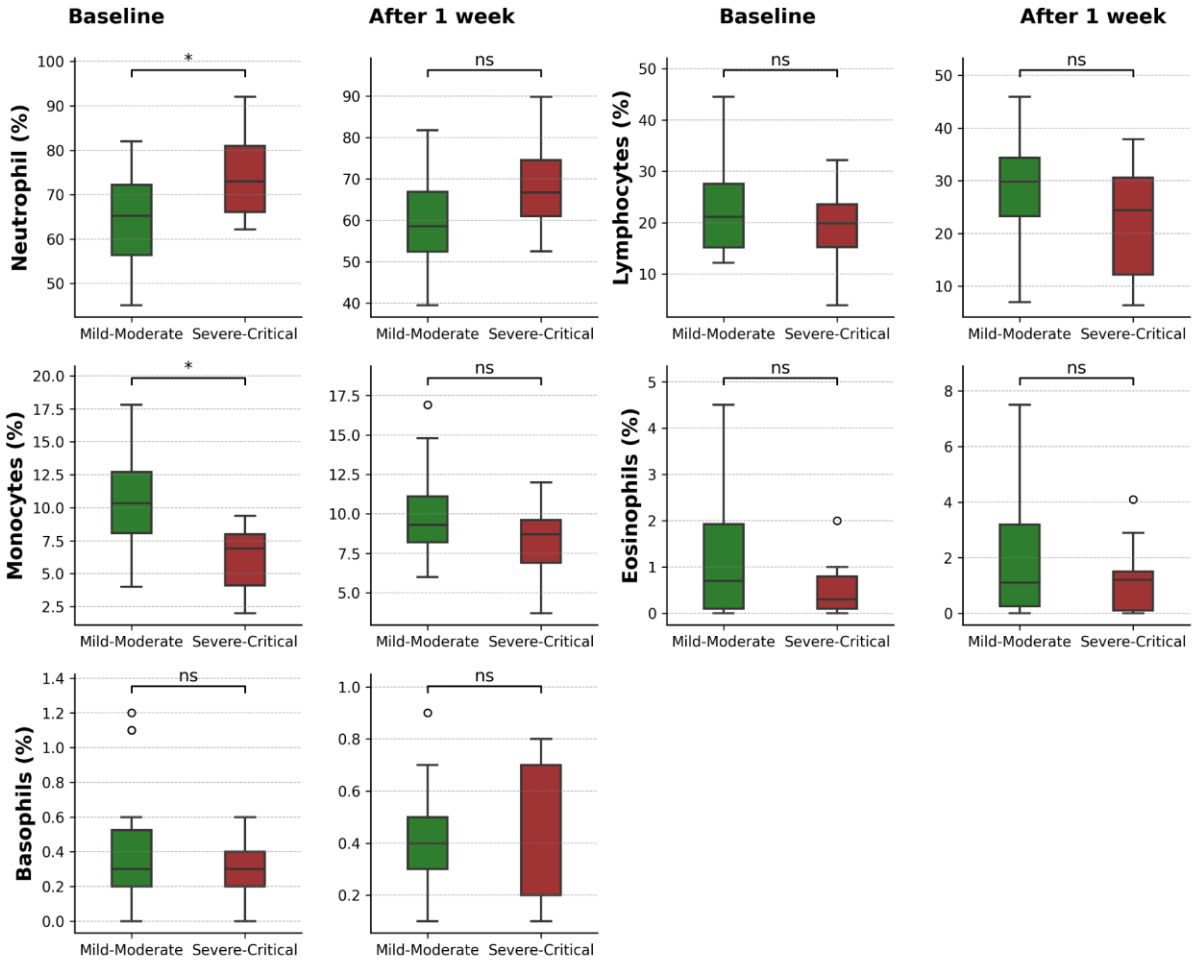


**Supplementary Fig 5. Baseline leukocyte proportion and its changes one week later in Mild-Moderate and Severe-Critical groups.** Boxplots show the proportions (%) of neutrophils, lymphocytes, monocytes, eosinophils, and basophils measured in whole blood at baseline (left panel) and after a week later (right panel) for patients with Mild–Moderate (green) and Severe–Critical (red) disease. Boxplots display the median (centre line), IQR (box limits), and 1.5×IQR (whiskers), and outliers (open circles). Statistical comparisons were conducted using the Wilcoxon rank-sum test. Statistical significance is denoted as follows: ns ($P \geq 0.05$), * ($P < 0.05$), ** ($P < 0.01$), *** ($P < 0.001$), **** ($P < 0.0001$).

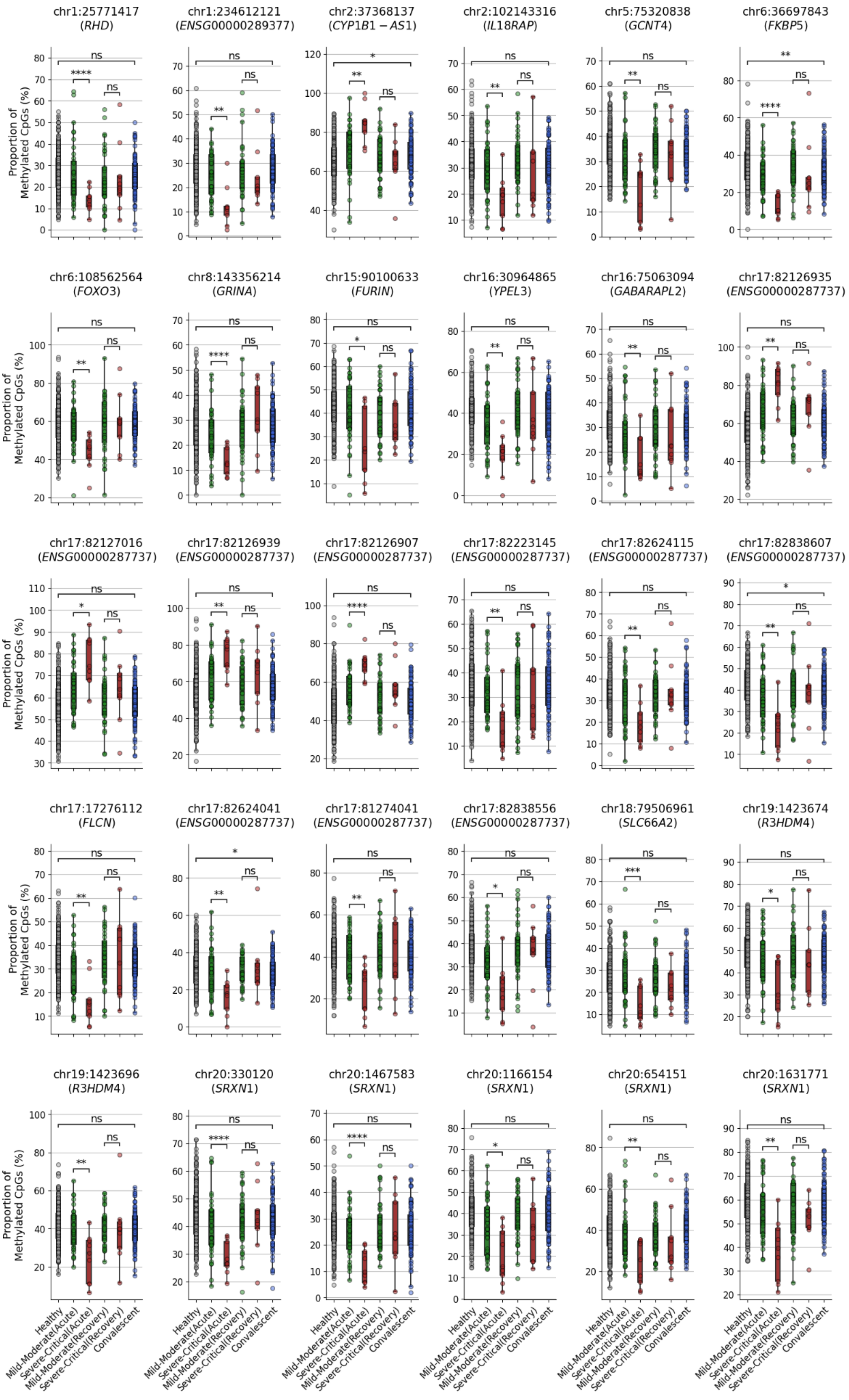
chr1:25771417 (RHD)
chr1:234612121 (ENSG00000289377)
chr2:37368137 (CYP1B1 − AS1)
chr2:102143316 (IL18RAP)
chr5:75320838 (GCNT4)
chr6:36697843 (FKBP5)
chr6:108562564 (FOXO3)
chr8:143356214 (GRINA)
chr15:90100633 (FURIN)
chr16:30964865 (YPEL3)
chr16:75063094 (GABARAPL2)
chr17:82126935 (ENSG00000287737)
chr17:82127016 (ENSG00000287737)
chr17:82126939 (ENSG00000287737)
chr17:82126907 (ENSG00000287737)
chr17:82223145 (ENSG00000287737)
chr17:82624115 (ENSG00000287737)
chr17:82838607 (ENSG00000287737)
chr17:17276112 (FLCN)
chr17:82624041 (ENSG00000287737)
chr17:81274041 (ENSG00000287737)
chr17:82838556 (ENSG00000287737)
chr18:79506961 (SLC66A2)
chr19:1423674 (R3HDM4)
chr19:1423696 (R3HDM4)
chr20:330120 (SRXN1)
chr20:1467583 (SRXN1)
chr20:1166154 (SRXN1)
chr20:654151 (SRXN1)
chr20:1631771 (SRXN1)
Proportion of Methylated CpGs (%)
Healthy
Mild-Moderate(Acute)
Severe-Critical(Acute)
Mild-Moderate(Recovery)
Severe-Critical(Recovery)
Convalescent

**Supplementary Fig 6. DNA Methylation Levels of 30 cis-eQTM CpGs Associated with COVID-19 Severity Across the Acute and Recovery Phases.** Methylation levels (% methylated CpGs) are shown for 30 *cis*-eQTM CpGs across clinical groups: Healthy controls (n = 310), patients with Mild-Moderate (n = 37) or Severe-Critical (n = 9) COVID-19 during the acute phase, and Convalescent individuals (n = 90). The x-axis indicates the clinical groups and infection periods. The y-axis represents the proportion of methylation at each CpG site. Boxplots display the median (centre line), IQR (box limits), 1.5×IQR (whiskers), and individual data points (closed circles). Statistical comparisons were performed using Welch's *t*-test. Statistical significance is denoted as follows: ns ($P \geq 0.05$), * ($P < 0.05$), ** ($P < 0.01$), *** ($P < 0.001$), **** ($P < 0.0001$).

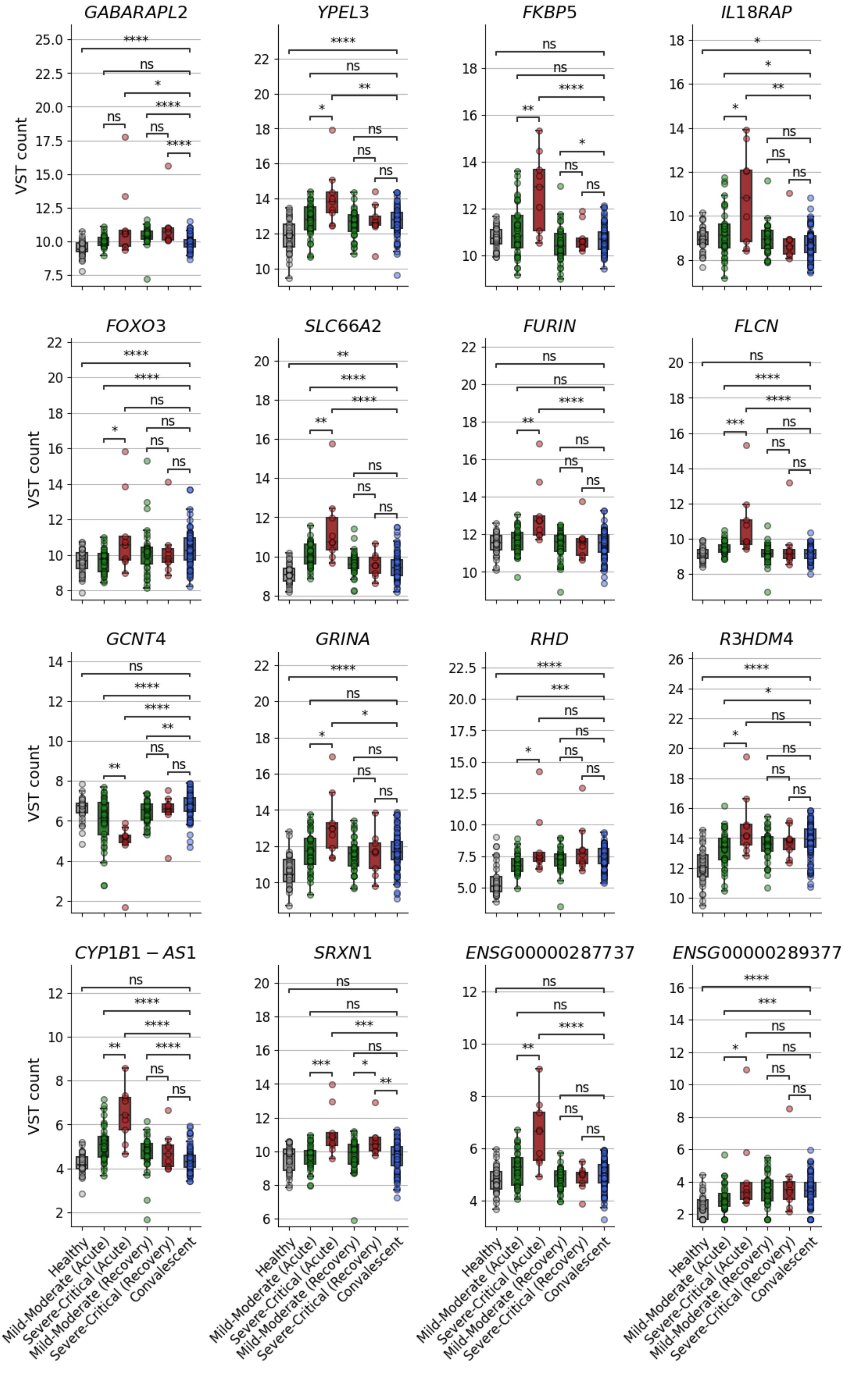
GABARAPL2
YPEL3
FKBP5
IL18RAP
FOXO3
SLC66A2
FURIN
FLCN
GCNT4
GRINA
RHD
R3HDM4
CYP1B1 − AS1
SRXN1
ENSG00000287737
ENSG00000289377
VST count
Healthy
Mild-Moderate (Acute)
Severe-Critical (Acute)
Mild-Moderate (Recovery)
Severe-Critical (Recovery)
Convalescent

**Supplementary Fig 7. Expression Levels of 16 *cis*-eQTM-Regulated Genes Associated with COVID-19 Severity Across the Acute and Recovery Phases.** Expression levels of 16 genes under cis-eQTM regulation are displayed across clinical groups: Healthy controls (n = 35), patients with Mild-Moderate (n = 36) or Severe-Critical (n = 9) COVID-19 during both the acute and recovery phase, and Convalescent individuals (n = 90). The x-axis indicates clinical groups and infection periods. The y-axis represents gene expression values, shown as variance-stabilizing transformed (VST) counts. Boxplots display the median (centre line), IQR (box limits), 1.5×IQR (whiskers), and individual data points (closed circles). Statistical comparisons were performed using the Wilcoxon rank-sum test. Statistical significance is reported as follows: ns ($P \geq 0.05$), * ($P < 0.05$), ** ($P < 0.01$), *** ($P < 0.001$), **** ($P < 0.0001$).

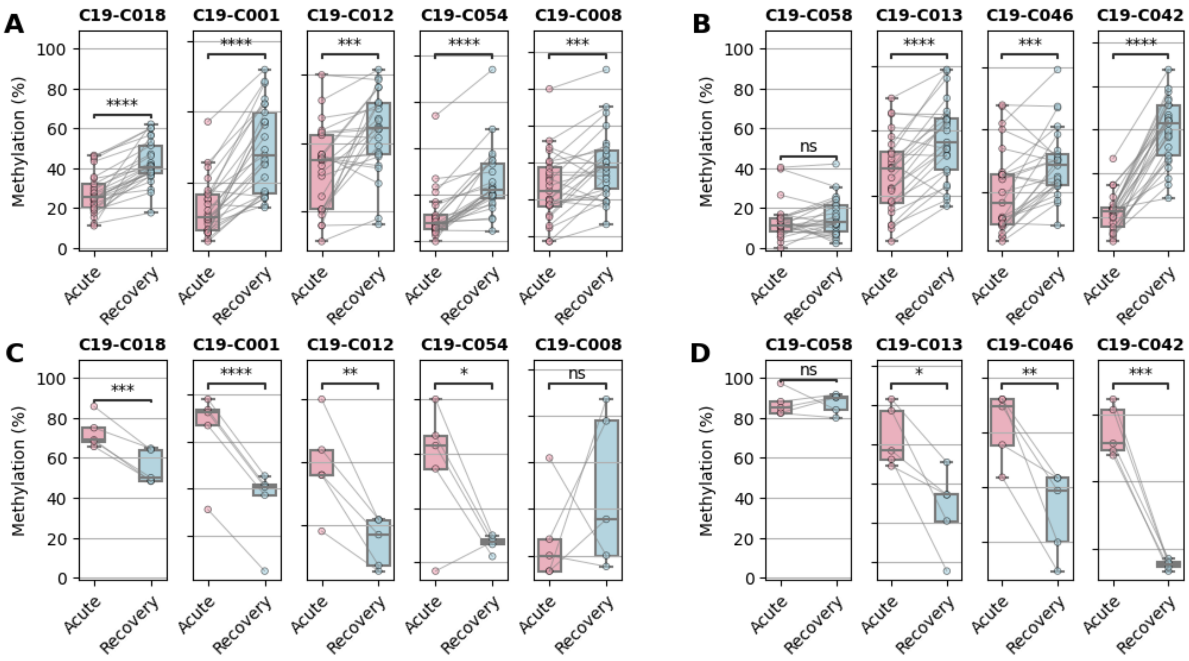


**Supplementary Fig 8. Paired longitudinal analysis of DNA methylation changes between acute and recovery phases in COVID-19 patients.** DNA methylation levels (β-values, %) at selected CpG sites are shown for matched whole blood samples collected from COVID-19 patients during acute illness and post-recovery. Panels **A** and **B** display CpG sites exhibiting hypomethylation during acute infection, while panels **C** and **D** show sites with hypermethylation, each stratified by clinical severity: **A, C)** severely ill and **B, D)** critically ill patients. Closed circles represent individual methylation β-values; grey lines denote paired measurements from the same patient across timepoints. Boxplots display the median (centre line), IQR (box limits), 1.5×IQR (whiskers), and individual data points within each group. Statistical differences between paired timepoints were evaluated using two-sided paired *t*-tests. Significance threshold is denoted as follows: ns ($P \geq 0.05$), * ($P < 0.05$), ** ($P < 0.01$), *** ($P < 0.001$), **** ($P < 0.0001$).

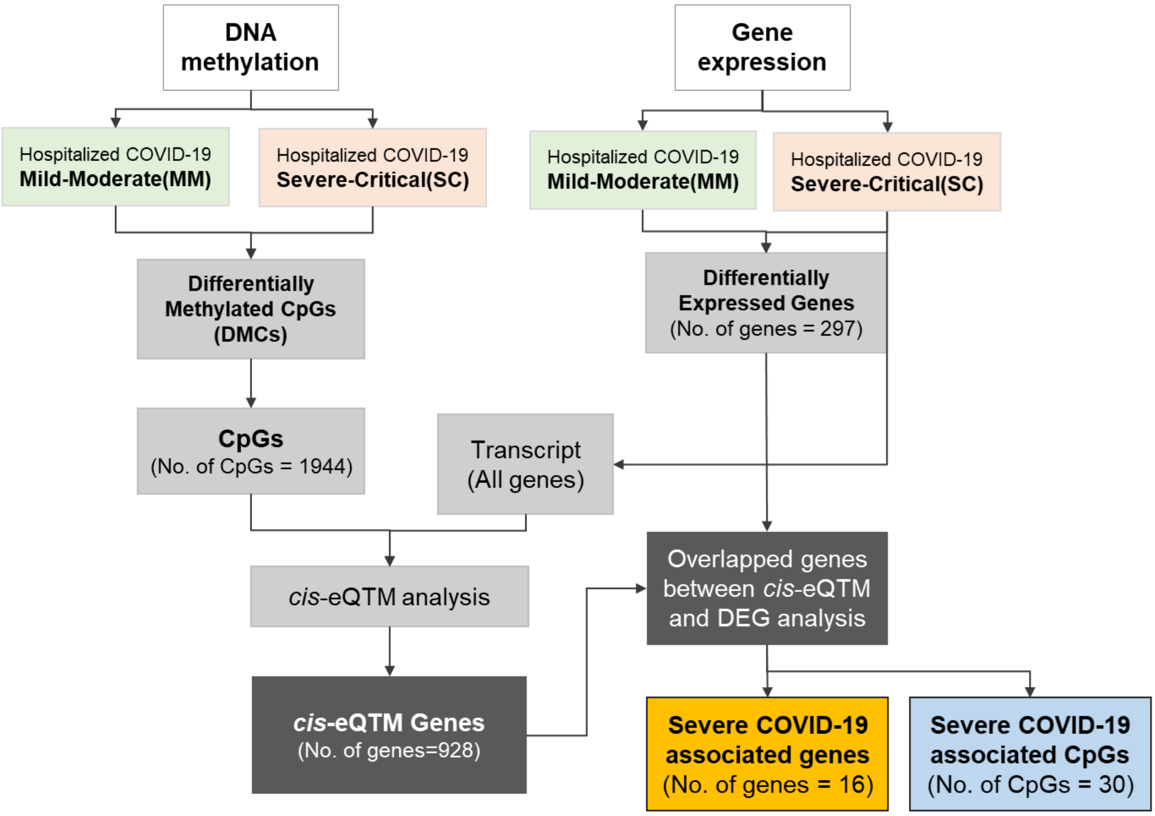


**Supplementary Fig 9. Overall process of COVID-19 *cis*-eQTM analysis.** The figure illustrates the workflow, including the discovery of differentially methylated CpGs and differentially expressed genes. Integration of methylation and gene expression data was achieved through the identification of *cis*-eQTM associations.

| Direction of methylation | Position | Gene symbol | Direction of mRNA expression | Correlation methylation with mRNA expression | | Median methylation differences(%) | Median gene expression differences (log2FC) |
|---|---|---|---|---|---|---|---|
| | | | | Spearman's ρ | P-value | | |
| Hyper-methylation | chr17:82126907 | *ENSG00000287737* | Up regulation | 0.571 | 0.006 | 13.517 | 1.515 |
| | chr17:82126935 | *ENSG00000287737* | Up regulation | 0.615 | 0.002 | 14.577 | 1.515 |
| | chr17:82126939 | *ENSG00000287737* | Up regulation | 0.526 | 0.015 | 11.995 | 1.515 |
| | chr17:82127016 | *ENSG00000287737* | Up regulation | 0.557 | 0.008 | 12.446 | 1.515 |
| | chr2:37368137 | *CYP1B1-AS1* | Up regulation | 0.512 | 0.020 | 14.037 | 1.536 |
| Hypo-methylation | chr1:25771417 | *RHD* | Up regulation | -0.530 | 0.014 | -11.900 | 2.037 |
| | chr1:234612121 | *ENSG00000289377* | Up regulation | -0.502 | 0.025 | -11.601 | 2.921 |
| | chr15:90100633 | *FURIN* | Up regulation | -0.602 | 0.003 | -14.306 | 1.596 |
| | chr16:30964865 | *YPEL3* | Up regulation | -0.544 | 0.010 | -15.843 | 1.466 |
| | chr16:75063094 | *GABARAPL2* | Up regulation | -0.569 | 0.006 | -10.430 | 2.249 |
| | chr17:17276112 | *FLCN* | Up regulation | -0.618 | 0.002 | -11.750 | 1.525 |
| | chr17:81274041 | *ENSG00000287737* | Up regulation | -0.567 | 0.006 | -16.428 | 1.515 |
| | chr17:82223145 | *ENSG00000287737* | Up regulation | -0.535 | 0.013 | -13.297 | 1.515 |
| | chr17:82624041 | *ENSG00000287737* | Up regulation | -0.631 | 0.001 | -12.341 | 1.515 |
| | chr17:82624115 | *ENSG00000287737* | Up regulation | -0.565 | 0.006 | -11.502 | 1.515 |
| | chr17:82838556 | *ENSG00000287737* | Up regulation | -0.675 | 0.000 | -11.950 | 1.515 |
| | chr17:82838607 | *ENSG00000287737* | Up regulation | -0.658 | 0.001 | -13.764 | 1.515 |
| | chr18:79506961 | *SLC66A2* | Up regulation | -0.607 | 0.002 | -13.415 | 1.562 |

| | | | | | | | |
|---|---|---|---|---|---|---|---|
| | chr19:1423674 | *R3HDM4* | Up regulation | -0.532 | 0.013 | -12.949 | 1.819 |
| | chr19:1423696 | *R3HDM4* | Up regulation | -0.557 | 0.008 | -14.906 | 1.819 |
| | chr2:102143316 | *IL18RAP* | Up regulation | -0.503 | 0.024 | -11.753 | 1.774 |
| | chr20:330120 | *SRXN1* | Up regulation | -0.582 | 0.004 | -11.078 | 1.517 |
| | chr20:654151 | *SRXN1* | Up regulation | -0.586 | 0.004 | -13.297 | 1.517 |
| | chr20:1166154 | *SRXN1* | Up regulation | -0.547 | 0.010 | -13.335 | 1.517 |
| | chr20:1467583 | *SRXN1* | Up regulation | -0.538 | 0.012 | -12.854 | 1.517 |
| | chr20:1631771 | *SRXN1* | Up regulation | -0.510 | 0.021 | -17.060 | 1.517 |
| | chr5:75320838 | *GCNT4* | Down regulation | 0.528 | 0.015 | -14.165 | -1.505 |
| | chr6:36697843 | *FKBP5* | Up regulation | -0.551 | 0.009 | -15.580 | 1.875 |
| | chr6:108562564 | *FOXO3* | Up regulation | -0.501 | 0.025 | -13.224 | 1.890 |
| | chr8:143356214 | *GRINA* | Up regulation | -0.546 | 0.010 | -10.716 | 1.490 |

**Supplementary Table 1. COVID-19 severity-associated genes identified through *cis*-eQTM analysis.** Columns indicate the direction of DNA methylation change (hypo- or hypermethylation), CpG site genomic position (hg38), associated gene symbol, direction of mRNA expression change (up- or downregulation), direction of correlation between DNA methylation and gene expression, Spearman's correlation coefficient (ρ), and associated *P*-value from *cis*-eQTM analysis adjusted for immune cell proportions.

| cis-eQTM CpG Site | Regulated Gene | Case | Control | Mean Difference | Test Statistics | *P*-value | FDR |
|---|---|---|---|---|---|---|---|
| chr2:102143316 | *IL18RAP* | Severe-Critical Acute | Healthy | -13.575 | -4.425 | 0.002 | 0.009 |
| chr2:102143316 | *IL18RAP* | Severe-Critical Acute | Mild-Moderate Acute | -11.566 | -3.301 | 0.005 | 0.01 |
| chr2:102143316 | *IL18RAP* | Healthy | Convalescent | 1.031 | 0.93 | 0.354 | 0.424 |
| chr2:102143316 | *IL18RAP* | Mild-Moderate Acute | Convalescent | -0.978 | -0.483 | 0.631 | 0.631 |
| chr2:102143316 | *IL18RAP* | Severe-Critical Acute | Convalescent | -12.544 | -3.947 | 0.003 | 0.009 |
| chr2:102143316 | *IL18RAP* | Mild-Moderate Acute | Healthy | -2.009 | -1.088 | 0.283 | 0.424 |
| chr5:75320838 | *GCNT4* | Severe-Critical Acute | Convalescent | -15.951 | -3.952 | 0.004 | 0.011 |
| chr5:75320838 | *GCNT4* | Healthy | Convalescent | 0.8 | 0.865 | 0.388 | 0.388 |
| chr5:75320838 | *GCNT4* | Mild-Moderate Acute | Convalescent | -2.561 | -1.362 | 0.179 | 0.215 |
| chr5:75320838 | *GCNT4* | Severe-Critical Acute | Healthy | -16.751 | -4.194 | 0.003 | 0.011 |
| chr5:75320838 | *GCNT4* | Mild-Moderate Acute | Healthy | -3.361 | -1.881 | 0.067 | 0.1 |
| chr5:75320838 | *GCNT4* | Severe-Critical Acute | Mild-Moderate Acute | -13.39 | -3.102 | 0.01 | 0.02 |
| chr6:36697843 | *FKBP5* | Severe_First | Mild_First | -15.916 | -5.915 | 5.76E-06 | 1.15E-05 |
| chr6:36697843 | *FKBP5* | Healthy | Convalescent | 3.258 | 2.624 | 0.01 | 0.011 |
| chr6:36697843 | *FKBP5* | Mild_First | Convalescent | -2.382 | -1.147 | 0.256 | 0.256 |
| chr6:36697843 | *FKBP5* | Severe_First | Convalescent | -18.298 | -7.987 | 2.17E-06 | 6.50E-06 |
| chr6:36697843 | *FKBP5* | Mild_First | Healthy | -5.641 | -3 | 0.004 | 0.007 |
| chr6:36697843 | *FKBP5* | Severe_First | Healthy | -21.556 | -10.198 | 1.90E-06 | 6.50E-06 |
| chr6:108562564 | *FOXO3* | Healthy | Convalescent | 0.273 | 0.241 | 0.81 | 0.81 |
| chr6:108562564 | *FOXO3* | Severe-Critical Acute | Convalescent | -14.679 | -4.665 | 9.60E-04 | 0.003 |
| chr6:108562564 | *FOXO3* | Mild-Moderate Acute | Healthy | -1.991 | -0.986 | 0.329 | 0.494 |
| chr6:108562564 | *FOXO3* | Severe-Critical Acute | Healthy | -14.953 | -4.891 | 9.60E-04 | 0.003 |

| | | | | | | | |
|---|---|---|---|---|---|---|---|
| chr6:108562564 | *FOXO3* | Severe-Critical Acute | Mild-Moderate Acute | -12.962 | -3.638 | 0.002 | 0.005 |
| chr6:108562564 | *FOXO3* | Mild-Moderate Acute | Convalescent | -1.717 | -0.798 | 0.428 | 0.514 |
| chr15:90100633 | *FURIN* | Severe-Critical Acute | Healthy | -17.069 | -3.256 | 0.011 | 0.045 |
| chr15:90100633 | *FURIN* | Healthy | Convalescent | 0.954 | 0.804 | 0.423 | 0.507 |
| chr15:90100633 | *FURIN* | Mild-Moderate Acute | Convalescent | -1.574 | -0.637 | 0.527 | 0.527 |
| chr15:90100633 | *FURIN* | Severe-Critical Acute | Convalescent | -16.115 | -3.032 | 0.015 | 0.045 |
| chr15:90100633 | *FURIN* | Mild-Moderate Acute | Healthy | -2.529 | -1.095 | 0.28 | 0.42 |
| chr15:90100633 | *FURIN* | Severe-Critical Acute | Mild-Moderate Acute | -14.541 | -2.563 | 0.026 | 0.052 |
| chr20:654151 | *SRXN1* | Mild-Moderate Acute | Convalescent | -3.281 | -1.337 | 0.187 | 0.28 |
| chr20:654151 | *SRXN1* | Severe-Critical Acute | Convalescent | -16.389 | -4.305 | 0.002 | 0.009 |
| chr20:654151 | *SRXN1* | Mild-Moderate Acute | Healthy | -2.141 | -0.93 | 0.358 | 0.358 |
| chr20:654151 | *SRXN1* | Severe-Critical Acute | Healthy | -15.249 | -4.109 | 0.003 | 0.009 |
| chr20:654151 | *SRXN1* | Severe-Critical Acute | Mild-Moderate Acute | -13.108 | -3.065 | 0.008 | 0.016 |
| chr20:654151 | *SRXN1* | Healthy | Convalescent | -1.139 | -0.93 | 0.354 | 0.358 |

**Supplementary Table 2. Summary table displaying pairwise comparisons of methylation levels at ten CpG sites identified through *cis*-eQTM analysis.** Columns indicate the genomic position of the CpG site (hg38), the regulated gene, the two comparison groups, the mean difference in methylation, test statistic (Welch's *t*-test), nominal *P*-value, and the false discovery rate (FDR).

| **Gene Symbol** | **Case** | **Control** | **Mean Difference** | **Test Statistics** | ***P*-value** | **FDR** |
|---|---|---|---|---|---|---|
| *ENSG00000115607.10_IL18RAP* | Severe-Critical Acute | Healthy | -1.966 | 2.342 | 0.019 | 0.038 |
| *ENSG00000115607.10_IL18RAP* | Severe-Critical Acute | Mild-Moderate Acute | -1.662 | 2.1 | 0.036 | 0.052 |
| *ENSG00000115607.10_IL18RAP* | Mild-Moderate Acute | Convalescent | -0.521 | 2.405 | 0.016 | 0.038 |
| *ENSG00000115607.10_IL18RAP* | Healthy | Convalescent | -0.217 | 2.023 | 0.043 | 0.052 |
| *ENSG00000115607.10_IL18RAP* | Mild-Moderate Acute | Healthy | -0.304 | 0.782 | 0.434 | 0.434 |
| *ENSG00000115607.10_IL18RAP* | Severe-Critical Acute | Convalescent | -2.183 | 3.073 | 0.002 | 0.013 |
| *ENSG00000176928.7_GCNT4* | Severe-Critical Acute | Mild-Moderate Acute | 1.117 | -2.611 | 0.009 | 0.014 |
| *ENSG00000176928.7_GCNT4* | Mild-Moderate Acute | Healthy | 0.63 | -2.277 | 0.023 | 0.027 |
| *ENSG00000176928.7_GCNT4* | Severe-Critical Acute | Convalescent | 1.92 | -4.725 | 2.30E-06 | 1.38E-05 |
| *ENSG00000176928.7_GCNT4* | Mild-Moderate Acute | Convalescent | 0.803 | -3.775 | 1.60E-04 | 3.20E-04 |
| *ENSG00000176928.7_GCNT4* | Healthy | Convalescent | 0.174 | -1.532 | 0.126 | 0.126 |
| *ENSG00000176928.7_GCNT4* | Severe-Critical Acute | Healthy | 1.746 | -4.263 | 2.02E-05 | 6.06E-05 |
| *ENSG00000096060.15_FKBP5* | Healthy | Convalescent | -0.035 | 0.441 | 0.659 | 6.59E-01 |
| *ENSG00000096060.15_FKBP5* | Mild-Moderate Acute | Convalescent | -0.357 | 1.286 | 0.198 | 0.297 |
| *ENSG00000096060.15_FKBP5* | Severe-Critical Acute | Convalescent | -1.992 | 3.557 | 3.75E-04 | 0.002 |
| *ENSG00000096060.15_FKBP5* | Mild-Moderate Acute | Healthy | -0.323 | 0.702 | 0.483 | 5.80E-01 |
| *ENSG00000096060.15_FKBP5* | Severe-Critical Acute | Healthy | -1.958 | 3.099 | 0.002 | 0.006 |
| *ENSG00000096060.15_FKBP5* | Severe-Critical Acute | Mild-Moderate Acute | -1.635 | 2.639 | 0.008 | 1.70E-02 |
| *ENSG00000118689.15_FOXO3* | Healthy | Convalescent | 0.702 | -3.761 | 1.69E-04 | 5.08E-04 |

| | | | | | | |
|---|---|---|---|---|---|---|
| *ENSG00000118689.15_FOXO3* | Mild-Moderate Acute | Convalescent | 0.664 | -3.802 | 1.44E-04 | 5.08E-04 |
| *ENSG00000118689.15_FOXO3* | Severe-Critical Acute | Convalescent | -0.777 | 0.348 | 0.728 | 0.809 |
| *ENSG00000118689.15_FOXO3* | Mild-Moderate Acute | Healthy | -0.039 | 0.242 | 0.809 | 0.809 |
| *ENSG00000118689.15_FOXO3* | Severe-Critical Acute | Mild-Moderate Acute | -1.441 | 1.958 | 0.05 | 0.075 |
| *ENSG00000118689.15_FOXO3* | Severe-Critical Acute | Healthy | -1.479 | 2.197 | 0.028 | 0.056 |
| *ENSG00000140564.13_FURIN* | Mild-Moderate Acute | Convalescent | -0.21 | 1.496 | 0.135 | 0.202 |
| *ENSG00000140564.13_FURIN* | Severe-Critical Acute | Mild-Moderate Acute | -1.458 | 3.121 | 0.002 | 0.004 |
| *ENSG00000140564.13_FURIN* | Severe-Critical Acute | Healthy | -1.627 | 3.797 | 1.47E-04 | 4.39E-04 |
| *ENSG00000140564.13_FURIN* | Mild-Moderate Acute | Healthy | -0.169 | 1.093 | 0.275 | 0.329 |
| *ENSG00000140564.13_FURIN* | Severe-Critical Acutee | Convalescent | -1.667 | 3.864 | 1.12E-04 | 4.39E-04 |
| *ENSG00000140564.13_FURIN* | Healthy | Convalescent | -0.041 | 0.195 | 0.845 | 0.845 |
| *ENSG00000271303.2_SRXN1* | Mild-Moderate Acute | Convalescent | -0.03 | -0.026 | 0.979 | 0.979 |
| *ENSG00000271303.2_SRXN1* | Healthy | Convalescent | 0.068 | -0.441 | 0.659 | 0.979 |
| *ENSG00000271303.2_SRXN1* | Severe-Critical Acute | Convalescent | -1.487 | 3.333 | 8.59E-04 | 0.002 |
| *ENSG00000271303.2_SRXN1* | Mild-Moderate Acute | Healthy | -0.098 | 0.23 | 0.818 | 0.979 |
| *ENSG00000271303.2_SRXN1* | Severe-Critical Acute | Healthy | -1.555 | 3.361 | 7.78E-04 | 0.002 |
| *ENSG00000271303.2_SRXN1* | Severe-Critical Acute | Mild-Moderate Acute | -1.456 | 3.32 | 9.01E-04 | 0.002 |

**Supplementary Table 3. Summary table displaying pairwise comparisons of expression levels at six genes identified through *cis*-eQTM analysis.** Columns report the gene symbol, comparison groups, mean difference in expression, test statistic (Wilcoxon rank-sum test), nominal *P*-value, and false discovery rate (FDR).

| Direction of methylation | Position | Gene symbol | Direction of mRNA expression | Correlation methylation with mRNA expression | |
|---|---|---|---|---|---|
| | | | | Spearman's ρ | *P*-value |
| Hypomethylation | chr6:36697843 | *FKBP5* | Up regulation | -0.551 | 0.00879 |

**Supplementary Table 4. A key gene associated with the severity of COVID-19 in the acute phase after adjustment for leukocyte composition.** Columns indicate the direction of DNA methylation change (hypo- or hypermethylation), CpG site genomic position (hg38), associated gene symbol, direction of mRNA expression change (up- or downregulation), direction of correlation between DNA methylation and gene expression, Spearman's correlation coefficient (ρ), and associated *P*-value from *cis*-eQTM analysis adjusted for immune cell proportions.

| Severity Level | Criteria |
|---|---|
| Mild | • Positive testing by virologic test (i.e., a nucleic acid amplification test of an antigen test)<br>• Symptoms of mild illness with COVID-19 that could include fever, cough, sore throat, malaise, headache, muscle pain, nausea, vomiting, diarrhea, and loss of taste or smell, without shortness of breath or dyspnea<br>• No clinical signs indicative of Moderate, Severe, or Critical Severity |
| Moderate | • Positive testing by virologic test (i.e., a nucleic acid amplification test of an antigen test)<br>• Symptoms of moderate illness with COVID-19, which could include any symptom of mild illness or shortness of breath with exertion<br>• Clinical signs suggestive of moderate illness with COVID-19, such as respiratory rate ≥ 20 breaths per minute, heart rate ≥ 90 beats per minute; with saturation of oxygen (SpO2) > 93% on room air at sea level<br>• No clinical signs indicative of Severe or Critical Illness Severity |
| Severe | • Positive testing by standard RT-PCR assay or an equivalent test<br>• Symptoms suggestive of severe systemic illness with COVID-19, which could include any symptom of moderate illness or shortness of breath at rest, or respiratory distress<br>• Clinical signs indicative of severe systemic illness with COVID-19, such as respiratory rate ≥ 30 per minute, heart rate ≥ 125 per minute, SpO2 ≤ 93% on room air at sea level or PaO2/FiO2 < 300<br>• No criteria for Critical Severity |
| Critical | • Positive testing by virologic test (i.e., a nucleic acid amplification test of an antigen test)<br>• Evidence of critical illness, defined by at least one of the following:<br>– Respiratory failure defined based on resource utilization requiring at least one of the following:<br>▪ Endotracheal intubation and mechanical ventilation, oxygen delivered by high flow nasal cannula (heated, humidified, oxygen delivered via reinforced nasal cannula at flow rates > 20 L/min with fraction of delivered oxygen ≥ 0.5), noninvasive positive pressure ventilation, ECMO, or clinical diagnosis of respiratory failure (i.e., clinical need for one of the preceding therapies, but preceding therapies not able to be administered in setting of resource limitation)<br>– Shock (defined by systolic blood pressure < 90 mm Hg, or diastolic blood pressure < 60 mm Hg or requiring vasopressors)<br>– Multi-organ dysfunction/failure |

**Supplementary Table 5. COVID-19 Severity Classification According to U.S. FDA Guidance (Feb 2021).** The classification of COVID-19 severity used in this study is based on the U.S. FDA's Guidance for Industry: COVID-19: Developing Drugs and Biological Products for Treatment or Prevention.

| CpGs | Healthy | Mild-Moderate | Severe-Critical | Convalescent |
|---|---|---|---|---|
| chr1:25771417 | 0.281 | 0.006 | 0.965 | 0.468 |
| chr1:234612121 | 0.382 | 0.382 | 0.382 | 0.726 |
| chr15:90100633 | 0.303 | 0.334 | 0.303 | 0.303 |
| chr16:30964865 | 0.877 | 0.877 | 0.877 | 0.281 |
| chr16:75063094 | **0.01** | **0.038** | **0.038** | 0.772 |
| chr17:17276112 | 0.606 | 0.665 | 0.606 | 0.872 |
| chr17:81274041 | 0.977 | 0.37 | 0.511 | 0.511 |
| chr17:82126907 | 0.085 | 0.085 | 0.653 | 0.653 |
| chr17:82126935 | 0.678 | 0.818 | 0.678 | 0.678 |
| chr17:82126939 | 0.703 | 0.703 | 0.703 | 0.703 |
| chr17:82127016 | 0.493 | 0.633 | 0.633 | 0.633 |
| chr17:82223145 | 0.061 | 0.061 | 0.572 | 0.643 |
| chr17:82624041 | 0.023 | 0.724 | 0.724 | 0.672 |
| chr17:82624115 | 0.024 | 0.636 | 0.636 | 0.612 |
| chr17:82838556 | 0.876 | 0.933 | 0.876 | 0.876 |
| chr17:82838607 | 0.762 | 0.762 | 0.762 | 0.762 |
| chr18:79506961 | **0.048** | 0.204 | 0.204 | 0.204 |
| chr19:1423674 | 0.386 | 0.838 | 0.357 | 0.386 |
| chr19:1423696 | 0.262 | 0.791 | 0.791 | 0.908 |
| chr2:37368137 | 0.647 | 0.324 | 0.647 | 0.647 |
| chr2:102143316 | 0.048 | 0.975 | 0.846 | 0.846 |
| chr20:330120 | 0.056 | 0.536 | 0.536 | 0.776 |
| chr20:654151 | **0.046** | **0.009** | 0.077 | 0.359 |
| chr20:1166154 | 0.809 | 0.646 | 0.646 | 0.646 |
| chr20:1467583 | 0.345 | 0.712 | 0.408 | 0.712 |
| chr20:1631771 | 0.746 | 0.746 | 0.746 | 0.746 |
| chr5:75320838 | 0.397 | 0.397 | 0.397 | 0.397 |
| chr6:36697843 | 0.377 | 0.961 | 0.332 | 0.332 |
| chr6:108562564 | 0.678 | 0.678 | 0.678 | 0.678 |
| chr8:143356214 | 0.571 | 0.849 | 0.571 | 0.854 |

**Supplementary Table 6. Summary table displaying significance of Shapiro-Wilk tests for 30 DMC-LOOs.** Each value represents an adjusted *P*-value of the normality test after Benjamini-Hochberg correction. Those values that have no evidence of normal distribution have been marked as bold font.

| Genes | Healthy | Mild-Moderate | Severe-Critical | Convalescent |
|---|---|---|---|---|
| ENSG00000287737.3_ENSG00000287737 | 0.997 | 0.949 | 0.949 | 0.802 |
| ENSG00000232973.14_CYP1B1-AS1 | 0.556 | 0.556 | 0.945 | **0.002** |
| ENSG00000118689.15_FOXO3 | 0.559 | 0.559 | 0.06 | **0.019** |
| ENSG00000154803.13_FLCN | 0.79 | 0.309 | **0.022** | 0.79 |
| ENSG00000115607.10_IL18RAP | 0.853 | 0.385 | 0.437 | 0.385 |
| ENSG00000187010.21_RHD | 0.002 | 0.347 | **0.003** | 0.132 |
| ENSG00000090238.12_YPEL3 | 0.663 | 0.663 | 0.138 | **0.034** |
| ENSG00000140564.13_FURIN | 0.608 | 0.608 | 0.055 | 0.364 |
| ENSG00000096060.15_FKBP5 | 0.634 | 0.318 | 0.634 | 0.318 |
| ENSG00000271303.2_SRXN1 | 0.059 | 0.254 | 0.114 | 0.059 |
| ENSG00000176928.7_GCNT4 | **0.028** | **0.02** | **0.002** | **0.02** |
| ENSG00000198858.10_R3HDM4 | 0.921 | 0.921 | 0.136 | **0.05** |
| ENSG00000122490.19_SLC66A2 | 0.864 | 0.864 | 0.063 | **0.009** |
| ENSG00000178719.17_GRINA | 0.741 | 0.793 | 0.524 | 0.741 |
| ENSG00000034713.8_GABARAPL2 | 0.539 | 0.539 | **0.011** | **0.032** |
| ENSG00000289377.2_ENSG00000289377 | **0.004** | **0.025** | **0.002** | **0.037** |

**Supplementary Table 7. Summary table displaying significance of Shapiro-Wilk tests for 16 DEG-LOOs.** Each value represents an adjusted *P*-value of the normality test after Benjamini-Hochberg correction. Those values that have no evidence of normal distribution have been marked as bold font.
